# Multidimensional Electroglottographic Characterization of Immediate Responses to Straw Phonation: A Pilot Methodological Study in Singers

**DOI:** 10.64898/2026.10.08.757648

**Authors:** Maya Little, Liran Oren

**Author notes:** Correspondence: Dr. L. Oren.

## Abstract

Semi-occluded vocal tract (SOVT) exercises are widely used in voice training and therapy, but immediate electroglottographic effects remain incompletely characterized. This pilot methodological study evaluated six female community singers using a no-straw baseline followed by Household metal, Metal VoiceStrawTM, plastic, and paper straw-phonation conditions. Thirty 1-s EGG recordings were analyzed using a standardized multidimensional pipeline including contact quotient (CQ), open quotient (OQ), waveform skewness, rise/fall timing, temporal skewness (s_t), normalized derivative asymmetry index (NDAI), average-cycle morphology, and spectral descriptors; participant-reported responses were summarized descriptively. Across all four post-exercise conditions, the EGG waveform showed a clear descriptive shift toward a steeper/faster closing transition relative to the slower opening transition, reflected by individual-level reductions in rise/fall ratio (4–5 of 6 participants per condition), higher s_t (three of four condition group means), and lower NDAI (all four conditions) than baseline. This closing-asymmetry pattern is consistent with a more rapid late-closing/contact transition, although EGG does not directly measure glottal airflow, maximum flow declination rate, or acoustic intensity. Post-SOVT spectra showed lower spectral centroid and more negative H2/H1, while low-order harmonic organization was largely preserved. Sensitivity analysis using 35%, 3/7 (42.9%), and 50% opening thresholds preserved the principal directional findings. These hypothesis-generating results support a multidimensional EGG framework for larger confirmatory studies.

## 1. Introduction

Semi-occluded vocal tract (SOVT) exercises are widely used in voice training, vocal rehabilitation, and singing pedagogy. These exercises partially constrict the vocal tract and include lip and tongue trills, humming, voiced fricatives, and phonation through tubes or straws [1]. Straw phonation is especially attractive because the oral constriction can be defined by the physical properties of the tube, making the exercise simple, reproducible, and adaptable to different vocal tasks [1–3]. For singers, SOVT exercises are commonly incorporated into warm-up and recovery routines with the aim of facilitating efficient phonation while limiting excessive vocal effort.

The physiological rationale for SOVT exercises is based on interaction between the glottal source and the acoustic and aerodynamic load imposed by the vocal tract. Semi-occlusion at the mouth increases supraglottal pressure and modifies vocal-tract impedance, thereby reducing the transglottal pressure difference and altering source–filter interaction [1]. Modeling and experimental studies indicate that these conditions can facilitate vocal-fold oscillation while modifying vibratory amplitude, collision behavior, and the pressure required to initiate or sustain phonation [1, 4]. Contemporary bioengineering studies likewise emphasize that phonation is a coupled fluid–structure–acoustic process in which vocal-tract resonances and supraglottal acoustics can feed back on vocal-fold vibration and vocal efficiency [5, 6].

A second mechanistic consideration is the temporal sharpness of vocal-fold closing. In glottal-source theory, acoustic excitation is strongly influenced by the rapid decline of transglottal airflow during the closing portion of the cycle. The maximum flow declination rate (MFDR), the most negative slope of the glottal-flow waveform, has been linked to vocal intensity and spectral excitation [7, 8]. Gauffin and Sundberg reported a high correlation between the negative peak of the differentiated flow glottogram and vowel sound-pressure level [7], while Titze showed theoretically that MFDR reflects contributions from both maximum area declination rate (MADR), a kinematic descriptor of glottal-area closing, and vocal-tract inertance [8]. This relationship is especially relevant to SOVT phonation because semi-occlusion can increase inertive reactance and source–tract interaction; modeling of tube phonation has explicitly used the MFDR/MADR relationship as an index of vocal economy [9]. Experimental flow studies further show that glottal-flow skewing, faster flow deceleration than acceleration, can increase glottal efficiency and can be augmented by intraglottal vortex dynamics during closing [10].

These established relationships motivate examining whether EGG cycle morphology contains a noninvasive signature of a more rapid closing/contact transition after straw exercise. EGG does not measure airflow, glottal area, MFDR, or MADR directly. Accordingly, the temporal skewness (s_t), rise/fall timing, and normalized derivative asymmetry index (NDAI) used here are interpreted as EGG-based markers of contact-waveform asymmetry. A shift toward a shorter EGG rise interval and a more abrupt contact transition is physiologically consistent with, but cannot by itself demonstrate, the higher MADR/MFDR behavior described in aerodynamic studies [8–10]. This distinction allows the present pilot to test a mechanistically important hypothesis without equating an electrical contact signal with a flow measurement.

SOVT exercises are not mechanically uniform interventions. Different semi-occlusions can generate substantially different aerodynamic loads. Maxfield et al. measured intraoral pressures produced by 13 SOVT gestures and demonstrated considerable differences among exercises [2], while electroglottographic measurements have shown that different SOVT tasks can produce different patterns of vocal-fold contact [11]. For flow-resistant tubes, diameter and length are established determinants of pressure–flow behavior, with diameter exerting a particularly strong influence on flow resistance [3]. Other work has examined phonation threshold pressure, voice-range behavior, and the combined effects of tube diameter, length, airflow, and water-immersion depth [4, 12, 13]. These findings indicate that the physical properties of an SOVT device are part of the intervention and should be defined explicitly.

Evidence concerning the effectiveness of SOVT exercises remains heterogeneous. Tube-based interventions have produced favorable outcomes in clinical populations and singers, including changes in aerodynamic, electroglottographic, acoustic, and perceptual measures [14]. However, systematic and scoping reviews have emphasized methodological heterogeneity, differences among exercise protocols, and the need for standardized approaches and broader samples [15, 16]. For a small pilot dataset, this literature supports emphasizing reproducible measurement and within-participant response patterns rather than treating the study as a definitive efficacy comparison.

A comparatively unresolved variable is straw material. Whereas tube diameter, length, airflow resistance, and immersion depth have received theoretical or experimental attention [2–4, 13], comparatively little evidence addresses whether the material from which a free-end straw is constructed influences short-term phonatory outcomes when geometry is maintained as nearly constant as practicable. Singers use devices made from stainless steel and other metals, plastic, paper, silicone, and related materials. Material could influence the user experience through wall stiffness, surface characteristics, thermal properties, tactile sensation, or subtle acoustic behavior; however, such effects should not be assumed. If pressure–flow behavior is governed predominantly by internal geometry and airflow [3], straws made from different materials but with closely matched dimensions may yield similar objective responses while still differing in comfort or perceived ease.

This distinction between objective vocal response and perceived benefit is particularly relevant for singers. The present study therefore combines electroglottography (EGG) with participant-reported outcomes. EGG provides a noninvasive measure related to changes in vocal-fold contact and has been used previously to characterize responses to SOVT exercises [11, 14]. In the present analysis, EGG-derived outcomes include contact quotient (CQ), open quotient (OQ), waveform-shape measures, temporal rise/fall asymmetry, derivative-based asymmetry, and normalized average-cycle morphology. Because EGG reflects electrical conductance associated with vocal-fold contact area rather than direct anatomical glottal closure, these quantities are treated as operational signal-derived measures rather than direct imaging-based measurements. This caution is consistent with the broader bioengineering literature, which treats vocal-fold vibration as a coupled biomechanical phenomenon whose measured behavior depends on the selected experimental representation and measurement method [6, 17].

Participant-reported outcomes were collected after testing to complement the EGG measures. A continuous microphone recording served only as procedural documentation and quality verification; no microphone-derived acoustic outcomes are analyzed in this manuscript.

Accordingly, this six-singer within-subject pilot evaluated immediate responses to four straw conditions using a reproducible EGG pipeline for contact, waveform morphology, temporal/derivative asymmetry, and spectral measures, with questionnaire responses as complementary perceptual outcomes. Particular attention was given to whether s_t, rise/fall timing, and NDAI showed a post-SOVT shift toward faster closing, thereby identifying candidate markers for future simultaneous EGG–airflow studies of MFDR and vocal efficiency.

## 2. Materials and Methods

### 2.1 Study Design and Scope of the Present Pilot Analysis

This study was conducted as a prospective, within-subject, repeated-measures investigation of semi-occluded vocal tract (SOVT) straw-phonation exercises in singers. The broader human-subjects protocol was designed to examine straw geometry and material using objective voice measurements and participant-reported outcomes. The present manuscript reports a pilot subset consisting of 30 electroglottographic (EGG) recordings from six participants together with the corresponding post-test questionnaire data. Each participant contributed five EGG recordings: a pre-exercise no-straw baseline and four recordings obtained after SOVT exercise using a Household metal straw, a Metal VoiceStrawTM, a plastic straw, or a paper straw. Each TDMS file contained 10,000 samples (1.00 s) from a single EGG channel sampled at 10 kHz.

Each participant served as their own baseline control and contributed data under every straw condition. Analyses focused on within-participant baseline-to-post-SOVT changes and descriptive differences among conditions. Quantitative outcomes were limited to EGG-derived measures and questionnaire responses; the microphone recording was used only for procedural documentation. Given n = 6, the study was descriptive and hypothesis-generating rather than inferential.

### 2.2 Participants, Recruitment, and Human-Subjects Protections

The participants in the tests were six female community singers, as approved by the IRB of the University of Cincinnati human-subjects protocol entitled “Effects of Straw Phonation Parameters on Vocal Efficiency.” Eligibility required self-reported good vocal health and willingness to complete the study procedures. Protocol exclusions included a history of vocal-fold pathology or laryngeal surgery, an ongoing voice disorder, recent upper-respiratory illness or allergy flare-up, current/recent voice therapy or voice-related medical treatment, and other factors considered likely to affect vocal quality. Screening was performed before study participation.

Participation was voluntary, and written informed consent was obtained before study-specific procedures. Participants could pause or stop testing at any time without penalty; no payment or financial incentive was provided.

The protocol classified the procedures as minimal risk. Tasks were brief and performed at comfortable pitch and loudness, with testing stopped if discomfort occurred. EGG and microphone measurements were non-invasive, and reusable study equipment was sanitized between participants.

Study data were coded with participant identifiers; consent/contact information was maintained separately, and research files were stored on secure University of Cincinnati systems accessible only to authorized personnel. Results are reported using de-identified or aggregate information.

### 2.3 Experimental Setup, Straw Conditions, and Test Procedure

The experimental station consisted of six principal components (Figure 1): (1) EGG neck strap/electrode assembly, (2) KAYPENTAX Model 6103 EGG controller, (3) microphone, (4) TASCAM US-2×2HR microphone power supply/audio interface, (5) NI cDAQ-9174 data-acquisition system (DAQ), and (6) acquisition computer. The EGG strap electrodes contacted the neck on either side of the larynx; the conditioned EGG signal was acquired through the DAQ. The microphone was recorded continuously as a procedural reference, not as an acoustic outcome.

**Figure 1.**
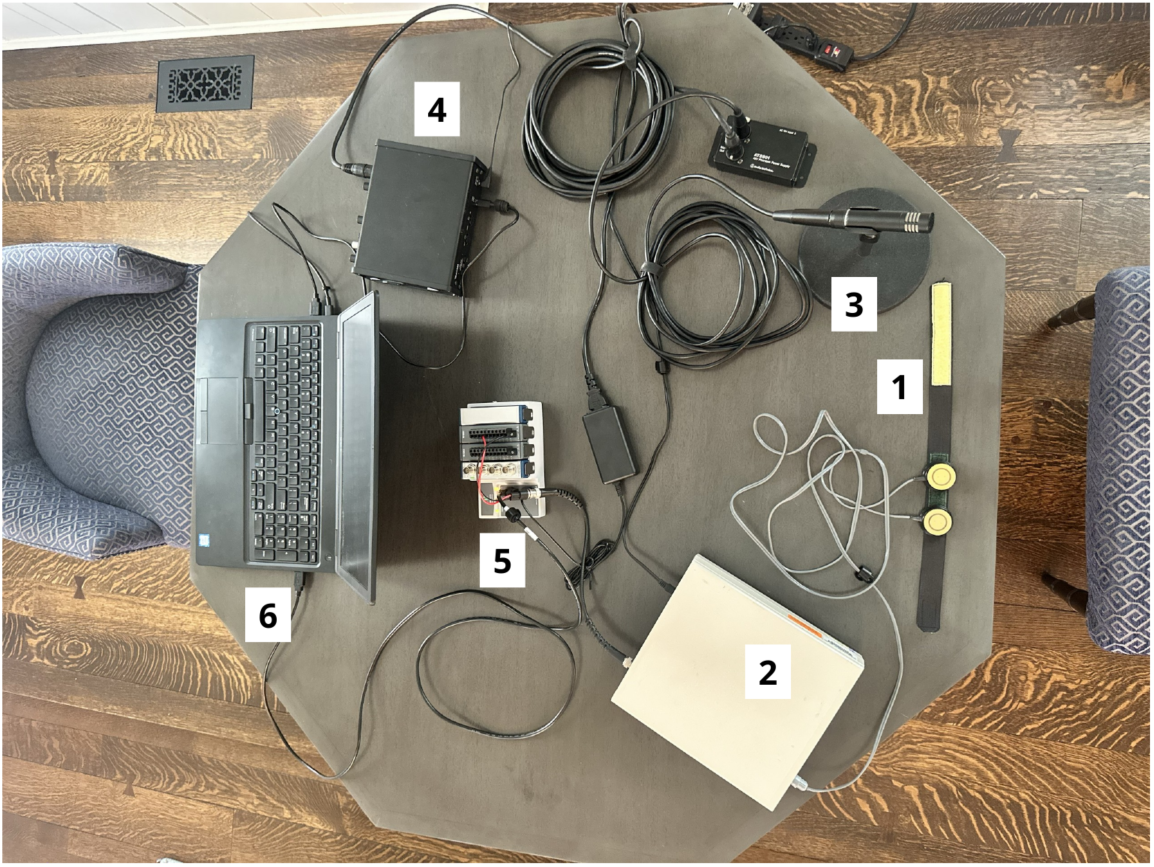
Experimental measurement setup. Numbered components are: (1) EGG neck strap/electrode assembly; (2) EGG controller, KAYPENTAX Model 6103; (3) microphone; (4) microphone power supply/audio interface, TASCAM US-2×2HR; (5) data-acquisition system (DAQ), NI cDAQ-9174; and (6) acquisition computer.

Four straw conditions were tested: a paper straw (length 179 mm, internal diameter [ID] 5.1 mm), a plastic straw (length 179 mm, ID 4.9 mm), a Metal VoiceStrawTM (length 178 mm, ID 5.7 mm), and a Household metal straw (length 180 mm, ID 4.4 mm) (Figure 2). Each participant completed the pre-exercise no-straw baseline and then the four SOVT conditions. The order of the four straw conditions was assigned across participants using a randomization scheme to balance order and carryover effects across the repeated-measures protocol. Between successive straw tests, participants completed a 10-min washout interval to reduce persistence of the preceding SOVT condition before the next test. The condition identity used in the analysis was taken from the filename/experimental record rather than inferred from file sequence. Table 1 shows the condition order for each participant.

**Figure 2.**
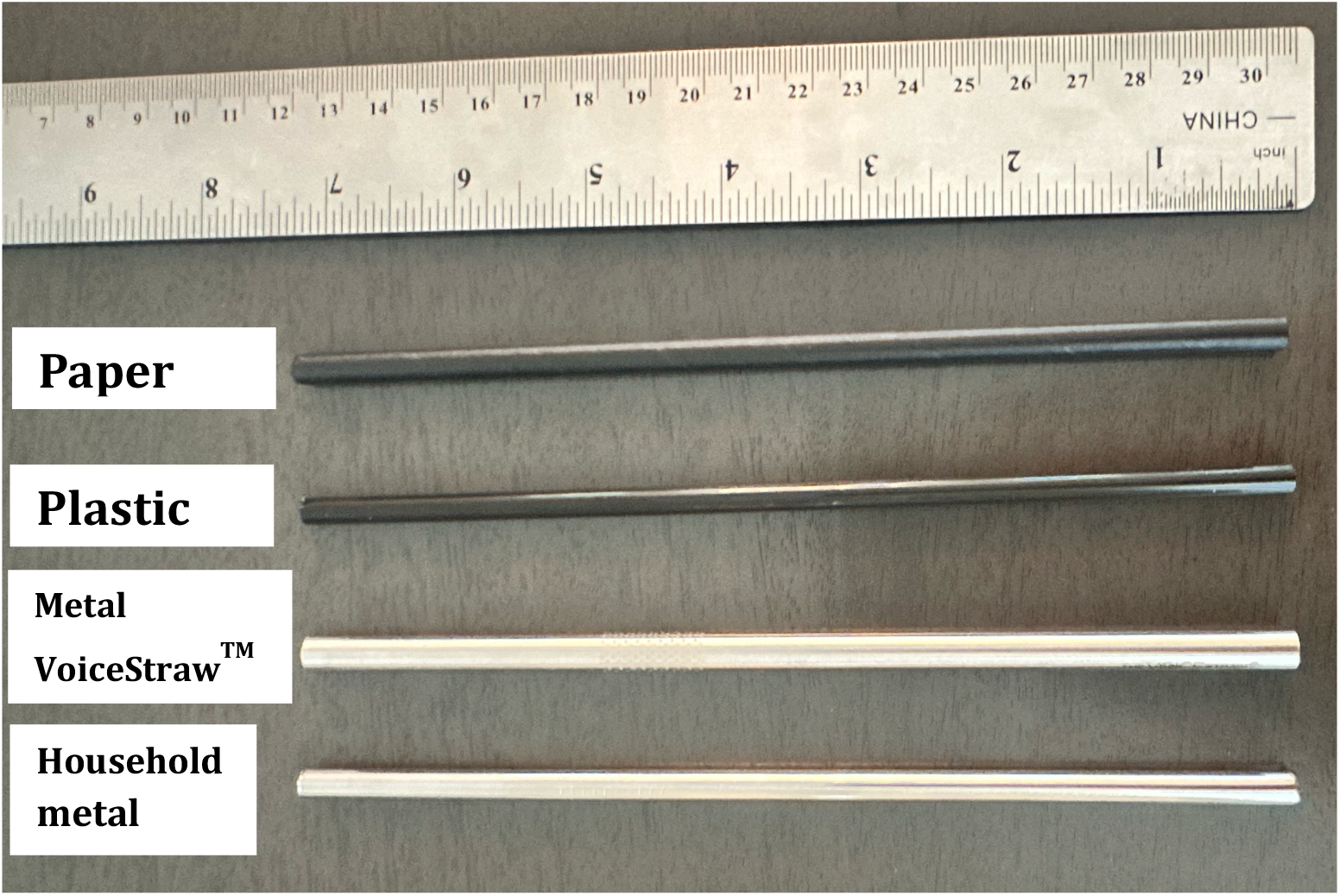
The four straw conditions evaluated in the study: paper (length 179 mm, ID 5.1 mm), plastic (length 179 mm, ID 4.9 mm), Metal VoiceStrawTM (length 178 mm, ID 5.7 mm), and Household metal (length 180 mm, ID 4.4 mm).

**Table 1.**
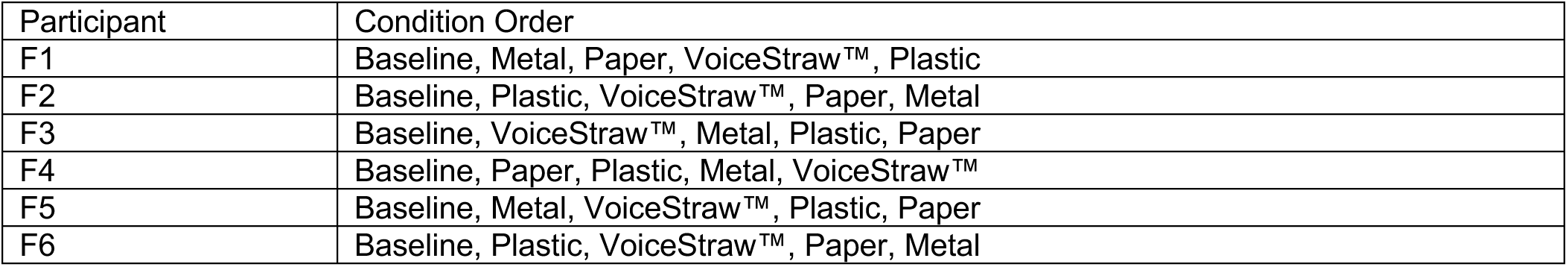
Participant condition order. Baseline was always first; the four straw conditions followed in the order shown.

| Participant | Condition Order |
| --- | --- |
| F1 | Baseline, Metal, Paper, VoiceStraw™, Plastic |
| F2 | Baseline, Plastic, VoiceStraw™, Paper, Metal |
| F3 | Baseline, VoiceStraw™, Metal, Plastic, Paper |
| F4 | Baseline, Paper, Plastic, Metal, VoiceStraw™ |
| F5 | Baseline, Metal, VoiceStraw™, Plastic, Paper |
| F6 | Baseline, Plastic, VoiceStraw™, Paper, Metal |

To reduce expectancy and visual-material bias, the straw was presented through a custom holder (Figure 3). The holder incorporated a mouthpiece connected to the test straw while the straw body and its material were concealed from direct view. Thus, the participant phonated through the mouthpiece without being shown which straw material was being tested.

**Figure 3.**
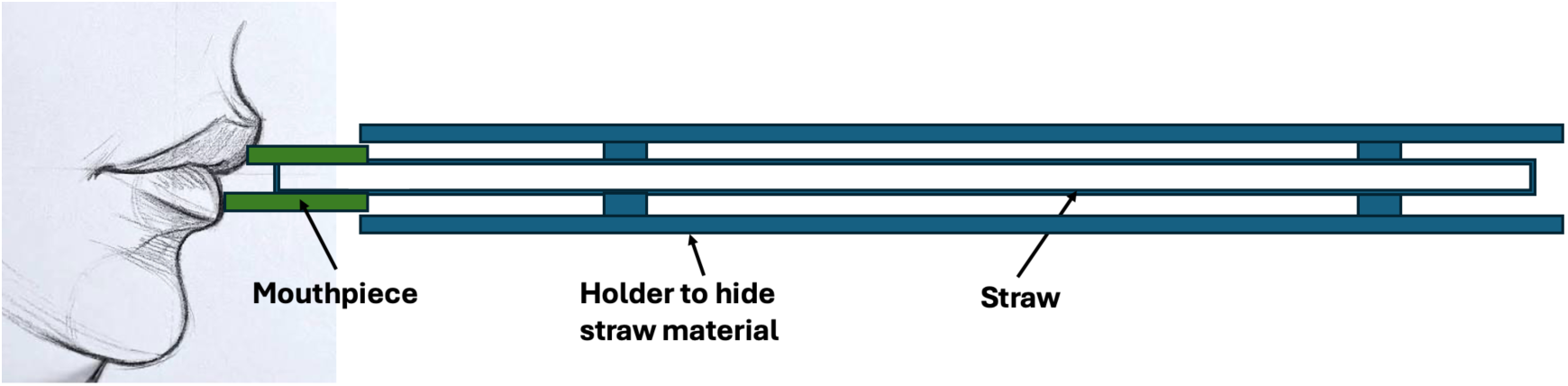
Custom straw-concealment holder. The participant phonated through the mouthpiece while the holder obscured direct visual identification of the straw material, reducing expectancy-related visual bias. 2.3a SOVT Exercise and Post-Exercise Recording Protocol

Participants performed straw phonations for 4 minutes. First, participants phonated at a comfortable self-selected pitch and loudness on a steady /oo/; then, vocal glides (degrees 1–5–1) and pentachordal scales (degrees 1–2– 3–4–5–4–3–2–1), both of which were sequentially transposed by a semitone up or down. Lastly, participants performed maximal-range vocal glissandi from the lowest sustainable pitch of their phonatory range to the highest, before returning to the baseline pitch. Each condition lasted for the duration of one minute. The exercise was performed in a seated position with the straw held at lip level. Immediately following the exercise, participants were asked to sing a steady /oo/ through the straw while a 1-s EGG recording was acquired. After this, the 10-minute vocal rest period began before repeating the straw phonations with a different straw. During this rest period, participants answered qualitative questions about the straws, as described in Section 2.10.

### 2.4 EGG Preprocessing

All 30 recordings were processed identically. Slow baseline drift was removed by subtracting a third-order Savitzky–Golay estimate computed with a 1001-sample (100.1-ms) window. The detrended EGG signal was then low-pass filtered at 1000 Hz with a fourth-order Butterworth filter applied in zero phase (forward–backward filtering). A differentiated electroglottographic signal (dEGG) was then calculated from the filtered EGG waveform using an 11-sample, third-order Savitzky–Golay differentiator. Mathematically, dEGG(t) = d[EGG(t)]/dt; therefore, dEGG expresses the instantaneous rate of change of the EGG conductance signal, not a second physiological recording. Figures 4 and 5 show the same illustrative baseline EGG recording and shows dEGG over 0.20–0.30 s. Sharp positive dEGG peaks identify the times at which EGG conductance rises most rapidly and were used operationally as rapid-contact (closure) landmarks. Negative excursions represent rapid decreases in EGG-measured contact associated with opening/separation. Differentiation makes these rapid transitions more prominent than in the slowly varying EGG waveform and therefore provides reproducible temporal landmarks for separating successive glottal cycles. The dEGG amplitude is expressed in signal units per second and should not be interpreted as contact area or force. Likewise, a positive dEGG peak is not direct imaging of the exact anatomical instant of closure; it marks the maximum rate of increase in EGG-measured contact under the present processing convention.

**Figure 4.**
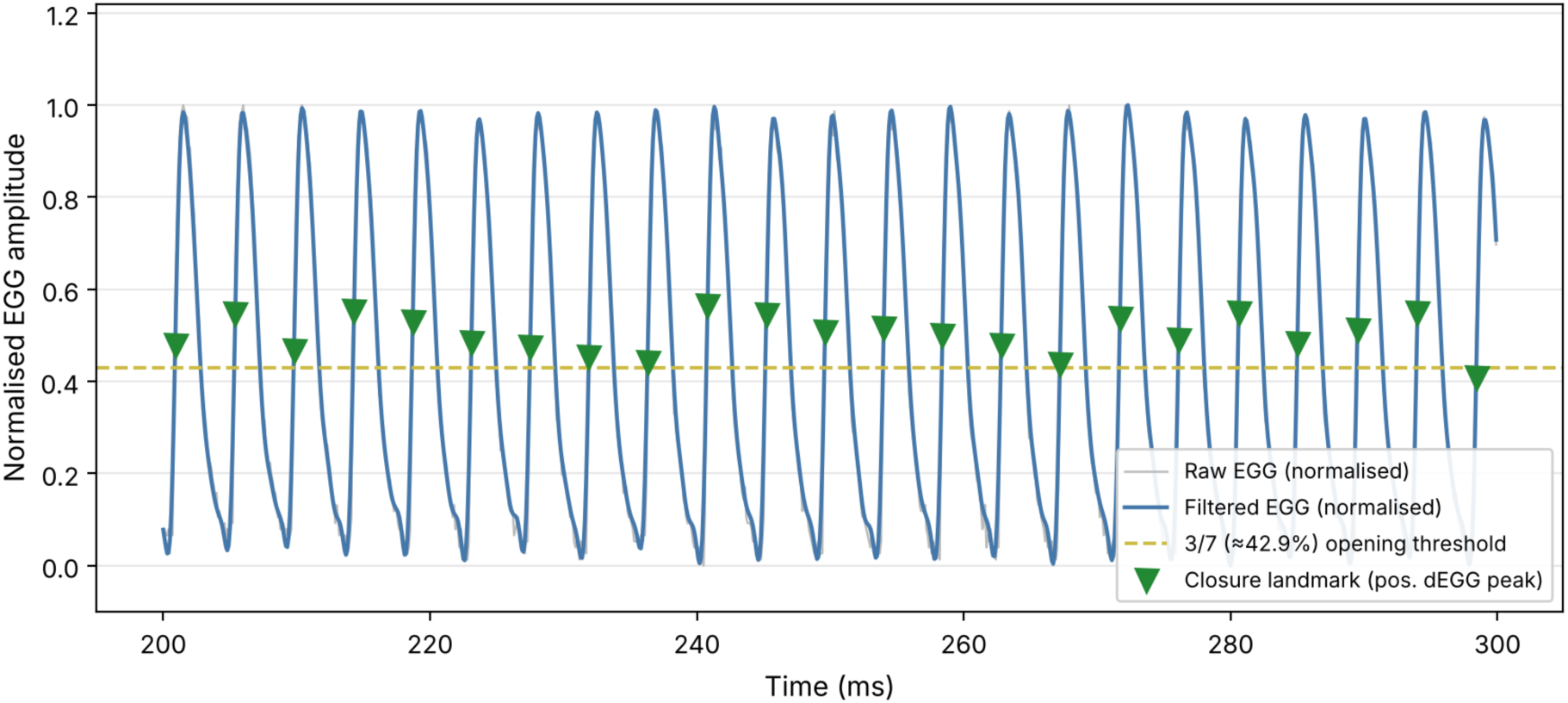
Illustrative baseline EGG cycle detection and CQ/OQ landmarks (0.20–0.30 s window, participant F6, baseline condition). Normalised raw (gray) and filtered (blue) EGG signals are shown together with the 3/7 (≈42.9%) opening-threshold line (dashed), positive-dEGG closure landmarks (green triangles), and 3/7-threshold opening landmarks (red circles). Shaded regions indicate the closed phase (green; used to compute CQ) and open phase (red; OQ) for one representative cycle.

**Figure 5.**
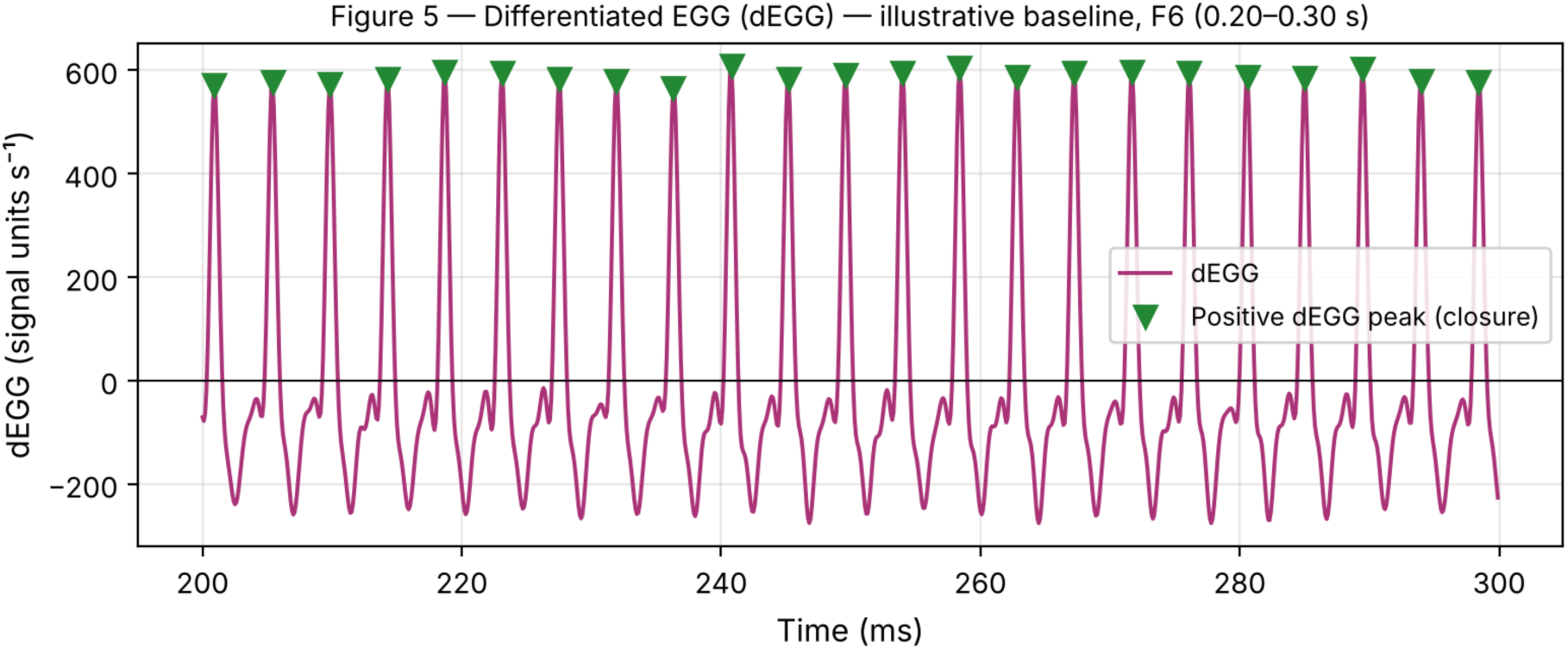
Differentiated EGG (dEGG) versus time for the same illustrative baseline recording shown in Figure 4 (participant F6, 0.20–0.30 s). Positive dEGG peaks (green triangles) mark the maximum rate of increase in EGG conductance and were used operationally as rapid-contact (closure) landmarks for glottal-cycle identification. dEGG amplitude is shown in signal units per second (s⁻¹).

### 2.5 Cycle Identification

Individual glottal cycles were defined by successive positive dEGG peaks, which were treated as rapid-contact (closure) landmarks (Figure 4). An initial period estimate was obtained from the dominant autocorrelation peak over lags corresponding to 70-500 Hz. Positive dEGG peaks were then detected with a minimum spacing of 0.65 times the estimated period and a minimum prominence of twice the median absolute deviation (MAD) of dEGG. Inter-peak intervals outside 60-550 Hz were rejected. Partial cycles at the beginning and end of each record were excluded. Across recordings, 147-352 complete cycles were retained (mean condition-level counts 246.5-270.5 cycles per 1-s recording).

### 2.6 Contact Quotient and Open Quotient

For each closure-to-closure cycle, the filtered EGG amplitude was normalized to the cycle-specific minimum and maximum. The primary opening/decontacting threshold was set a priori at 3/7 (42.9%) of the cycle amplitude above the minimum. This choice follows the Howard hybrid EGG approach, in which the relatively distinct positive dEGG peak is used to mark the rapid contacting event while a fixed amplitude crossing of the EGG waveform is used for the more gradual and often less distinctly marked decontacting/opening event [18, 19]. Howard’s 3/7 criterion has been used in subsequent EGG work for this purpose [19]. The opening landmark was the first downward threshold crossing after the within-cycle EGG maximum. Contact quotient (CQ) was calculated as the elapsed time from the positive dEGG closure landmark to this crossing divided by the closure-to-closure period; open quotient (OQ) was defined as 1 - CQ. Because EGG reflects contact-related conductance rather than glottal area or airflow, the threshold crossing was treated as an operational EGG decontacting landmark rather than an exact anatomical opening instant. Cycles yielding implausible CQ values below 0.05 or above 0.95 were excluded.

Opening-threshold sensitivity analysis. To determine whether the principal condition patterns depended on the 3/7 criterion, all 30 recordings were reanalyzed using 35%, 3/7 (42.9%), and 50% normalized opening thresholds while holding preprocessing, dEGG closure landmarks, and accepted cycle boundaries fixed. The 35% and 50% criteria were selected because both have precedent in EGG criterion-level methods [20]. For each threshold, CQ/OQ, fall time, rise/fall ratio, temporal skewness (s_t), and NDAI were recomputed. Rise time itself is independent of the opening threshold because its endpoint is the within-cycle EGG maximum. The purpose of this analysis was not to select the threshold that maximized a condition difference, but to test whether the direction of the baseline-to-straw findings was robust to reasonable alternative opening criteria.

[Sensitivity analysis results reported in Section 3.1a.]

### 2.7 Waveform Shape, Rise/Fall Timing, and Asymmetry Measures

Waveform skewness was computed from the filtered EGG sample values within each accepted cycle. Three complementary descriptors were used to characterize rise/fall asymmetry. Note that rise and fall intervals span only the contact phase of each cycle (approximately CQ·T), not the full glottal cycle; accordingly, these measures characterize contact-phase timing asymmetry rather than whole-cycle temporal asymmetry. Rise time was defined as the interval from the positive dEGG closure landmark to the within-cycle EGG maximum, and fall time as the interval from that maximum to the 3/7 opening-threshold crossing. Their ratio (rise time/fall time) was retained as an intuitive landmark-based supporting measure: a value of 1 denotes equal durations, whereas values below 1 indicate a shorter rise interval than fall interval. Temporal skewness (s_t) and the Normalized Derivative Asymmetry Index (NDAI), defined below, were treated as the principal complementary asymmetry outcomes because they characterize whole-cycle temporal asymmetry and derivative-shape asymmetry, respectively. Derivative skewness was calculated separately over the rise and fall intervals for use in NDAI and was interpreted as a waveform-shape descriptor rather than a direct measure of glottal force.

The two principal dimensionless asymmetry measures were calculated from the same accepted cycles. Temporal skewness (s_t) quantified the relative difference between the fall and rise intervals as s_t = (T_fall - T_rise)/(T_fall + T_rise), where T_rise and T_fall are the cycle-specific rise and fall durations defined above. Thus, s_t = 0 denotes equal durations, positive values indicate a longer fall interval, and negative values indicate a longer rise interval. The Normalized Derivative Asymmetry Index (NDAI) quantified the relative imbalance between the magnitudes of derivative skewness during the fall and rise portions: NDAI = (|S_d,fall| - |S_d,rise|)/(|S_d,fall| + |S_d,rise|), where S_d,rise and S_d,fall are the skewness values of dEGG samples in the rise and fall portions of each cycle. Absolute values were used because rise-and fall-segment derivative skewness can have opposite signs. Both indices are bounded from −1 to +1. Cycle-level values were averaged within each recording and then summarized across the six participants for each condition. A separate bounded timing index was not retained because (T_rise - T_fall)/(T_rise + T_fall) is algebraically identical to −s_t and would therefore provide no independent information. Consequently, s_t is a monotonic transform of the rise/fall ratio (s_t = (1−r)/(1+r)); their concordance is a consistency check, not independent corroboration. Post-straw rise times of approximately 0.55–0.62 ms correspond to 5–6 samples at 10 kHz, which is shorter than the 11-sample Savitzky– Golay differentiator window; absolute timing values should therefore be interpreted with caution. NDAI is computed from the skewness of approximately 6 (rise) and 11 (fall) samples per cycle; bootstrap analysis across cycles within a recording is recommended in future work to assess its stability.

For mechanistic interpretation, the three descriptors were considered together rather than as competing measures. The rise/fall ratio provides a readily interpretable landmark-based check on transition timing; s_t provides the principal normalized temporal-asymmetry measure; and NDAI provides a complementary derivative-based description that emphasizes differences in the shapes and rates of the rising and falling contact transitions. Concordant directional changes across these measures were considered stronger descriptive evidence of altered cycle morphology than a change in any single metric. Under the present landmark convention, an increase in s_t accompanied by a shorter rise time and lower rise/fall ratio indicates that the EGG waveform reaches its contact maximum more rapidly after the positive dEGG closure landmark, relative to the subsequent fall/opening interval. These variables were prespecified as EGG contact-transition descriptors, not as direct estimates of MFDR or MADR. MFDR requires a glottal-airflow waveform (typically obtained with inverse-filtered oral airflow or another validated aerodynamic method), and MADR requires a glottal-area measure. The EGG asymmetry metrics were therefore used to identify patterns that could be tested against simultaneous airflow/area measurements in a larger study [8–10].

### 2.8 Average-Cycle Construction

Each accepted cycle was amplitude-normalized from 0 to 1 and resampled by linear interpolation to 201 equally spaced points over one closure-to-closure period. Cycles were first averaged within each recording to obtain 30 participant-by-condition mean cycles. Grand-average cycles were then calculated across the six participant means for each condition, thereby weighting participants equally rather than weighting recordings by their number of detected cycles. Recording-level values were likewise obtained by averaging cycle-level measurements, and group values are reported as mean ± SD across the six participants. Because this was a six-participant pilot dataset, interpretation emphasizes within-participant patterns, descriptive direction and magnitude of change, and methodological robustness. No population-level inferential hypothesis testing was used to determine differences among conditions. The resulting condition-specific grand-average waveforms and their morphological comparison are presented in Section 3.6 (Figure 10).

### 2.9 EGG Spectral and Time–Frequency Analysis

The preprocessed 1-s EGG recordings were additionally characterized in the frequency domain. Power spectral density was estimated with Welch’s method using Hann windows, 2048-sample segments (204.8 ms), 50% overlap, and a 4096-point FFT; quantitative spectral metrics were calculated over 20-1000 Hz, consistent with the retained EGG bandwidth. For visualization, the spectra were displayed from 0 to 1000 Hz so that the low-frequency boundary and harmonic series were readily apparent. The corresponding spectrograms used 512-sample Hann windows (51.2 ms), 448-sample overlap, and a 2048-point FFT and were likewise displayed from 0 to 1000 Hz. Because absolute EGG amplitude depends strongly on electrode contact and placement, spectral displays were normalized and interpreted as waveform/contact-spectrum measures rather than calibrated acoustic spectra.

Fundamental frequency was estimated from the dominant autocorrelation period. Harmonic ratios H2/H1 and H3/H1 were expressed in decibels as 10log10(PH2/PH1) and 10log10(PH3/PH1), respectively. Spectral centroid was calculated as the power-weighted mean frequency from 20 to 1000 Hz. Spectral slope was obtained by least-squares regression of spectral level (dB) against frequency (kHz) from 100 to 1000 Hz. For each participant, post-SOVT values were also normalized to that participant’s baseline: centroid was expressed as percentage change, whereas slope and harmonic ratios were expressed as arithmetic dB differences. These measures describe the spectrum of the EGG contact waveform and should not be equated directly with microphone-derived acoustic spectral measures.

### 2.10 Perceptual and Self-Reported Measures

Participants completed a post-condition questionnaire during the protocol. For each straw, the form asked whether singing felt easier, harder, or unchanged after the exercise and whether the voice felt different (yes/no), with space to describe any perceived change. After all four conditions, participants ranked the four straws from 1 (most effective) to 4 (least effective) for improving vocal efficiency, identified the straw that made singing feel easiest overall, and indicated which straw they would be most likely to use for future warm-ups. These responses were summarized descriptively because of the six-participant pilot sample. Comparative rankings were tabulated by first-choice frequency and mean rank; open-ended comments were summarized qualitatively. No Likert composite score was calculated.

### 2.11 Procedural Audio Recording and Quality Documentation

A continuous microphone recording was retained only to verify task delivery, session chronology, and the presence of the brief test phonations during EGG acquisition, and to distinguish measurement segments from warm-up, rest, questionnaire, and conversational periods. It was not calibrated or analyzed for acoustic outcomes.

### 2.12 Statistical Analysis

Because this was a six-participant pilot methodological study, analyses were descriptive and hypothesis-generating rather than population-level inferential. For EGG outcomes, recording-level cycle summaries were reduced to participant-condition means and then summarized across participants as mean ± SD. Within-participant baseline-to-post-condition differences, baseline-normalized changes, ranges, and graphical comparisons were used where informative. No population-level hypothesis tests, post-hoc significance tests, or multiplicity-adjusted inferential comparisons were used to determine differences among straw conditions.

Questionnaire responses were summarized descriptively using condition-specific easier/harder/no-change and voice-difference responses, comparative rank order, first-choice counts, and open-ended comments.

## 3. Results

### 3.1 Cycle detection and fundamental frequency

Cycle detection yielded analyzable periodic EGG data in all 30 recordings. The number of retained cycles per recording ranged from 152 to 361. Mean F0 was 280.3 ± 56.7 Hz at baseline (corrected; see Section 2.5 QC note), 260.9 ± 56.0 Hz after the Household metal straw, 275.4 ± 29.6 Hz after the Metal VoiceStrawTM, 271.8 ± 51.1 Hz after the plastic straw, and 262.0 ± 52.9 Hz after the paper straw. Because participants were not constrained to identical post-exercise F0 values, waveform-shape comparisons should be interpreted together with these pitch differences.

#### 3.1a Opening-Threshold Sensitivity Analysis

As a methodological robustness check, the alternative thresholds (35%, 3/7 [42.9%], and 50%) changed absolute criterion-dependent values as expected but preserved the principal descriptive directions. Across all three thresholds, all four straw-condition mean CQs remained above baseline; all four straw conditions retained lower mean rise/fall ratios and higher mean s_t than baseline; and mean NDAI remained below baseline. Baseline CQ was 0.412, 0.378, and 0.350 and baseline s_t was 0.327, 0.272, and 0.217 at 35%, 3/7, and 50%, respectively. These checks support retaining 3/7 as the prespecified primary criterion while recognizing that absolute EGG timing values are threshold dependent.

### 3.2 Contact and open quotients

Mean baseline CQ was 0.378 ± 0.112 (OQ 0.622 ± 0.112). Mean CQ increased to 0.408 ± 0.091 for the Household metal straw, 0.436 ± 0.085 for the Metal VoiceStrawTM, 0.421 ± 0.066 for the plastic straw, and 0.405 ± 0.094 for the paper straw. Corresponding mean OQ values were 0.592, 0.564, 0.579, and 0.595. Thus, all four post-exercise condition means showed a numerically higher CQ than baseline, with the largest group-mean difference for the Metal VoiceStrawTM (+0.058). Individual responses were heterogeneous; for example, some participants showed lower CQ after particular straw conditions despite the positive group-mean changes. The CQ and OQ condition means are summarized graphically in Figure 6.

**Figure 6.**
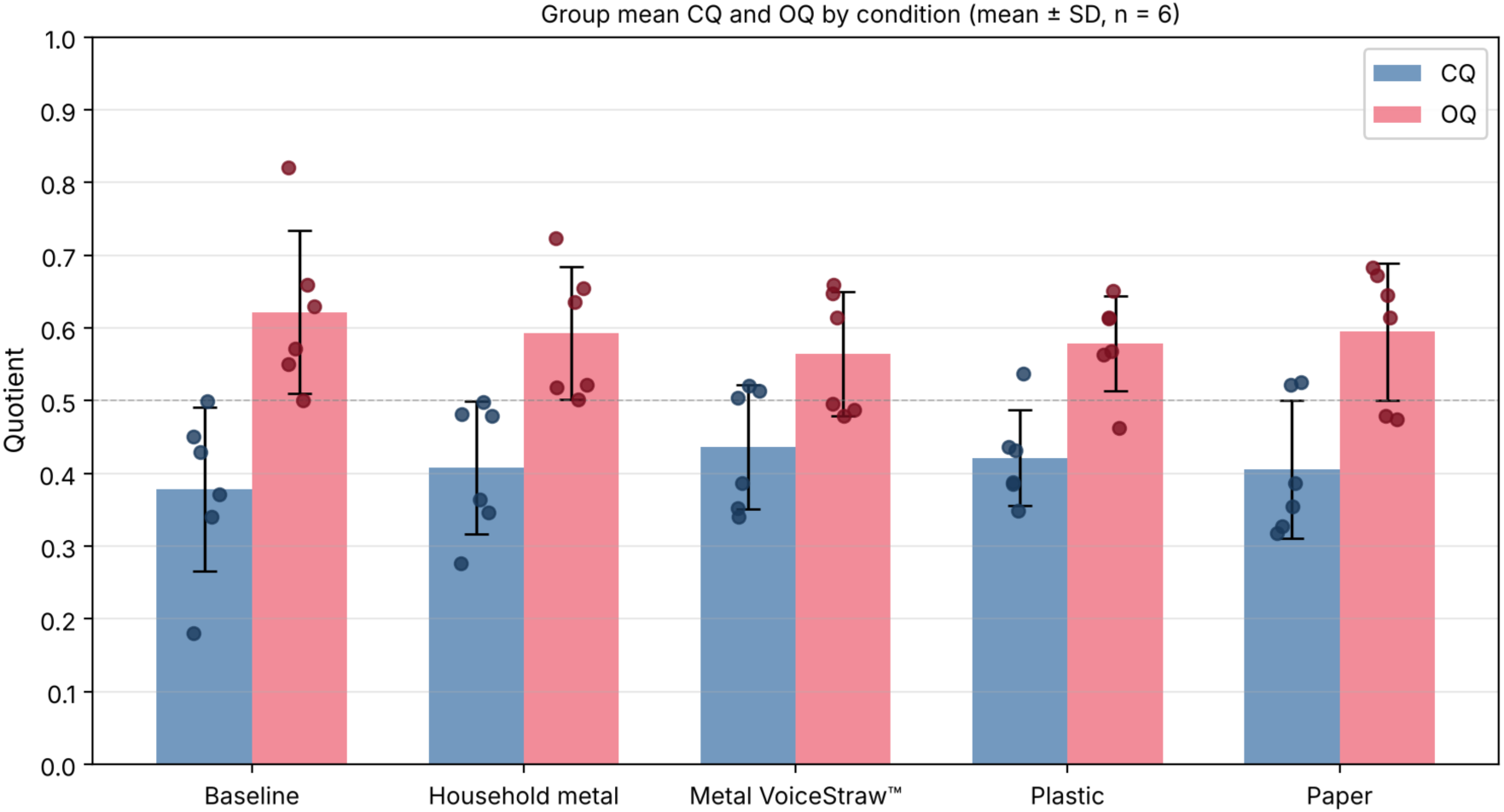
Group mean contact quotient (CQ; blue) and open quotient (OQ; red) across the five conditions. Error bars show ±1 SD across the six participants; filled points show individual participant values. Note that CQ + OQ = 1 by definition.

### 3.3 Waveform skewness

Mean EGG waveform skewness was 0.394 ± 0.208 at baseline. Post-exercise means were 0.355 ± 0.316 for the Household metal straw, 0.198 ± 0.236 for the Metal VoiceStrawTM, 0.291 ± 0.233 for the plastic straw, and 0.333 ± 0.291 for the paper straw. All four post-exercise group means were numerically lower than baseline, with the largest decrease for the Metal VoiceStrawTM. These condition means are shown in Figure 7.

**Figure 7.**
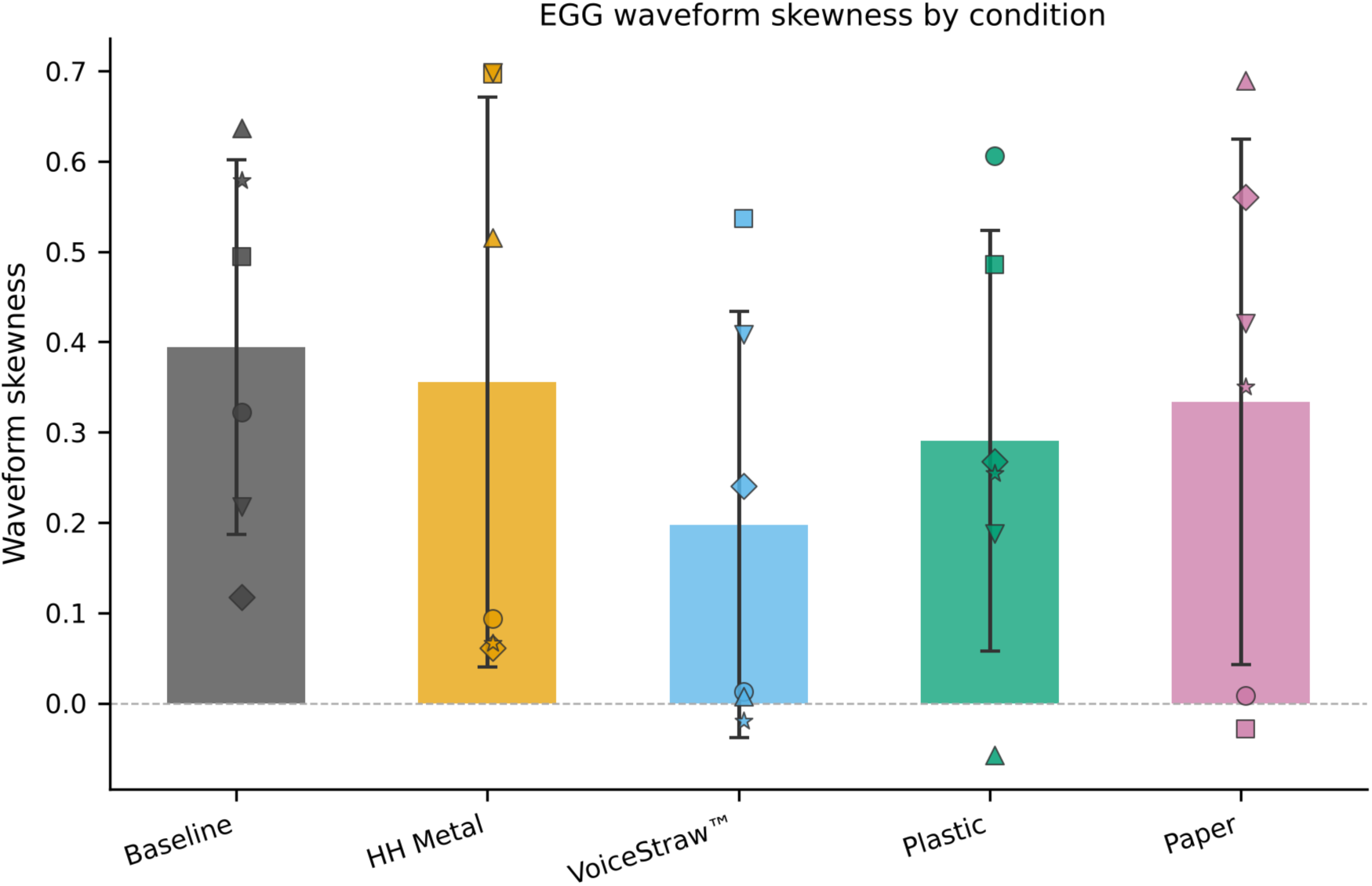
EGG waveform skewness by condition. Bars show group means ± SD across six participants; filled points show individual participant values.

### 3.4 Rise/fall timing asymmetry

The baseline mean rise time was 0.476 ± 0.089 ms and mean fall time was 0.870 ± 0.298 ms, producing a mean rise/fall ratio of 0.588 ± 0.135 (all six participants). Within-participant direction counts for rise/fall decrease (post < baseline): Household metal 4/6, VoiceStraw 5/6, plastic 4/6, paper 3/6. These counts indicate a modest directional tendency rather than a uniform shift. Because the rise/fall ratio is retained as an intuitive supporting descriptor rather than a primary asymmetry outcome, the timing findings are interpreted as hypothesis-generating rather than confirmatory. Baseline variability was substantial, so the pattern should be regarded as descriptive.

### 3.5 Temporal skewness and normalized derivative asymmetry

The principal temporal-asymmetry outcome, temporal skewness (s_t), was positive on average in every condition. Mean ± SD s_t was 0.272 ± 0.103 at baseline, 0.287 ± 0.178 after the Household metal straw, 0.295 ± 0.176 after the Metal VoiceStrawTM, 0.331 ± 0.122 after the plastic straw, and 0.241 ± 0.168 after the paper straw. Accordingly, three of the four post-exercise condition means (Household metal, Metal VoiceStrawTM, and plastic) showed greater temporal asymmetry toward a longer fall interval than the baseline mean; the paper condition mean (0.241) fell marginally below baseline (0.272). The largest mean s_t was observed after the plastic straw. Participant-level responses remained heterogeneous; these group means are therefore interpreted descriptively. The participant-level and group-mean temporal-skewness values are displayed in Figure 8.

**Figure 8.**
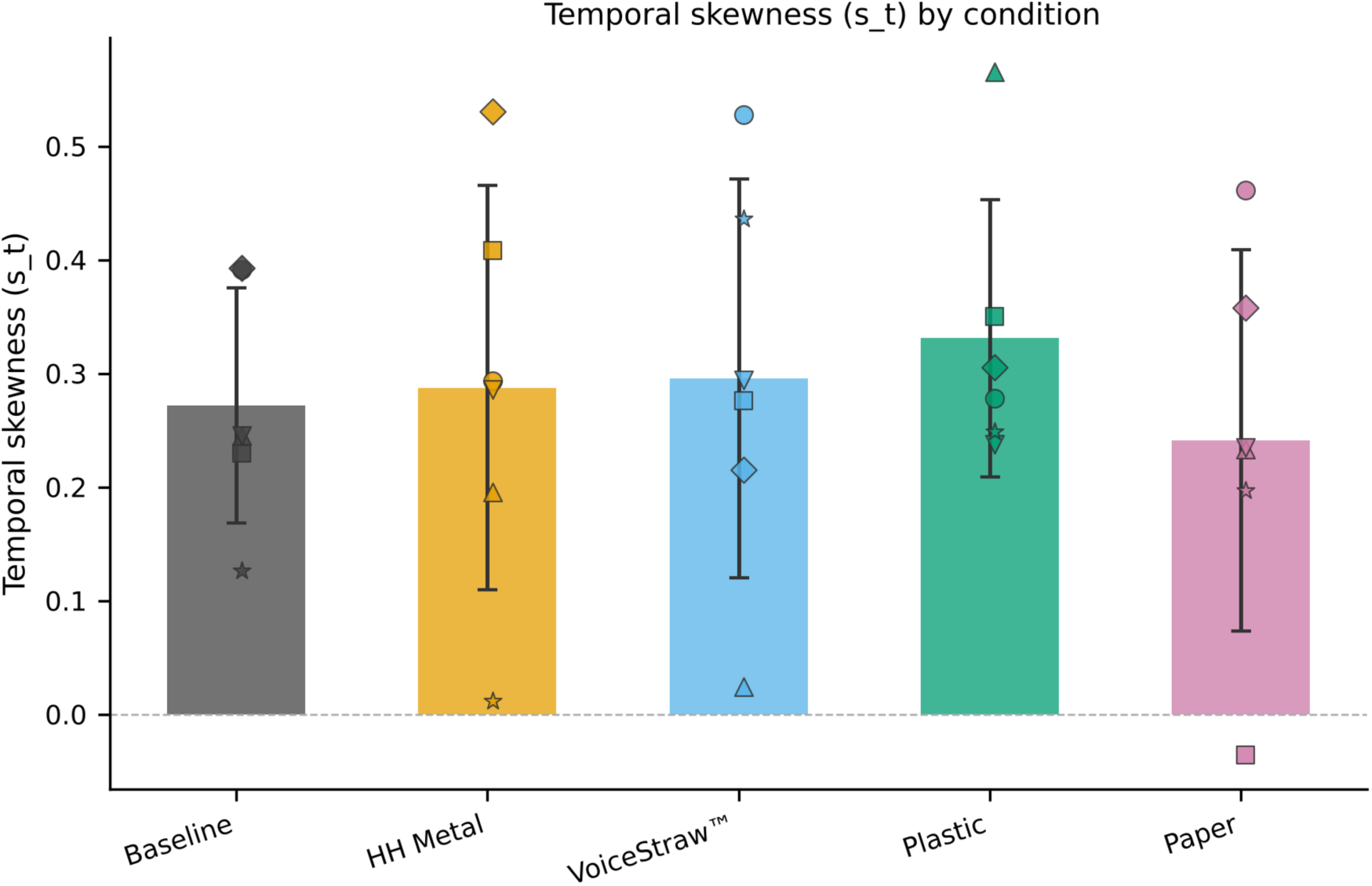
Temporal skewness (s_t) across baseline and the four post-exercise straw conditions. Bars show group means ± SD across six participants; filled points show individual participant values. Positive s_t indicates a longer fall (EGG closure) interval than rise (EGG opening) interval.

The complementary derivative-based outcome, NDAI, showed a concordant descriptive change. Mean ± SD NDAI was 0.368 ± 0.240 at baseline, 0.270 ± 0.182 after the Household metal straw, 0.253 ± 0.173 after the Metal VoiceStrawTM, 0.330 ± 0.154 after the plastic straw, and 0.348 ± 0.207 after the paper straw. Thus, the group mean NDAI was lower than baseline after each straw exercise, with the largest descriptive reductions for Metal VoiceStrawTM and Household metal. These differences are presented descriptively and are not interpreted as population-level condition effects. The corresponding NDAI values are displayed in Figure 9.

**Figure 9.**
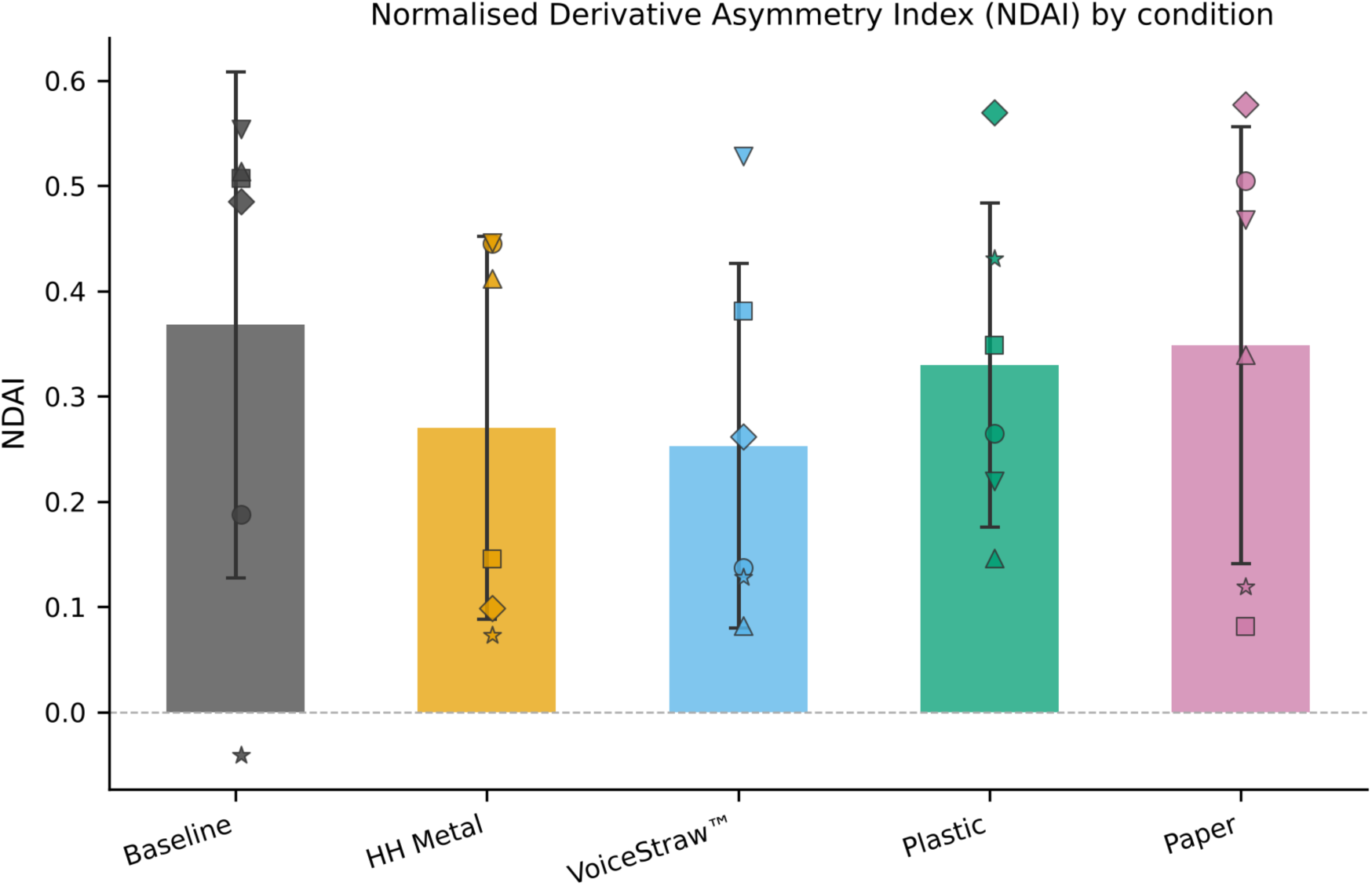
Normalized Derivative Asymmetry Index (NDAI) across baseline and the four post-exercise straw conditions. Bars show group means ± SD across six participants; filled points show individual participant values. Positive NDAI indicates that the dEGG closing slope is steeper (larger absolute value) than the opening slope.

Considered jointly, the timing measures showed a coherent descriptive shift toward a faster EGG contact transition after straw phonation. Mean absolute rise time was 0.476 ms at baseline and 0.554–0.614 ms across the four straw conditions; fall time was 0.870 ms at baseline and 1.042–1.191 ms across conditions. Because F0 was lower in most straw conditions, both rise and fall durations increased in absolute terms, so the relative asymmetry metrics (rf_ratio, s_t, NDAI) are the primary interpretive focus. The baseline rise/fall ratio was 0.588 ± 0.135; five of six participants showed a lower rf_ratio after the Metal VoiceStrawTM condition and four of six after Household metal and plastic. Group-mean rf_ratios for Household metal (0.600), Metal VoiceStrawTM (0.628), and paper (0.664) were modestly above baseline (0.588), while plastic (0.562) was below, reflecting high participant-level variability pulling the metal-straw and paper means upward. The s_t pattern was concordant: three of four post-exercise group means exceeded baseline (0.272), with Household metal (0.287), Metal VoiceStrawTM (0.295), and plastic (0.331) all higher; the paper mean (0.241) was slightly below baseline. NDAI decreased from 0.368 at baseline to 0.253–0.348 after the straw conditions; all four post-exercise group means were below baseline. Because these are EGG contact-waveform measures, the data should be described as consistent with a more rapid late-closing/contact transition, not as direct evidence that MFDR itself increased. Numerical summaries of s_t and NDAI are provided in Table 2.

**Table 2.** Temporal skewness and NDAI summary. Values are mean ± SD across six participants. Positive s_t indicates a longer fall interval than rise interval. Positive NDAI indicates greater derivative-skewness magnitude in the fall portion than the rise portion.

| Condition | Temporal skewness, $s_t$ | NDAI |
| --- | --- | --- |
| Baseline | $0.272 \pm 0.103$ | $0.368 \pm 0.240$ |
| Household metal | $0.287 \pm 0.178$ | $0.270 \pm 0.182$ |
| Metal VoiceStrawTM | $0.295 \pm 0.176$ | $0.253 \pm 0.173$ |
| Plastic | $0.331 \pm 0.122$ | $0.330 \pm 0.154$ |
| Paper | $0.241 \pm 0.168$ | $0.348 \pm 0.207$ |

Values are mean ± SD across six participants. Positive $s_t$ indicates a longer fall interval than rise interval. Positive NDAI indicates greater derivative-skewness magnitude in the fall portion than the rise portion.

### 3.6 Average-Cycle Morphology

The grand-average normalized cycles showed visible condition-related changes in waveform morphology. Although the curves were broadly similar in overall periodic shape, they differed in the timing and curvature of the high-contact portion and the subsequent decline. These descriptive waveform differences provide a visual complement to the quantitative asymmetry measures reported above and should be interpreted as exploratory in this six-participant pilot dataset.

**Figure 10.**
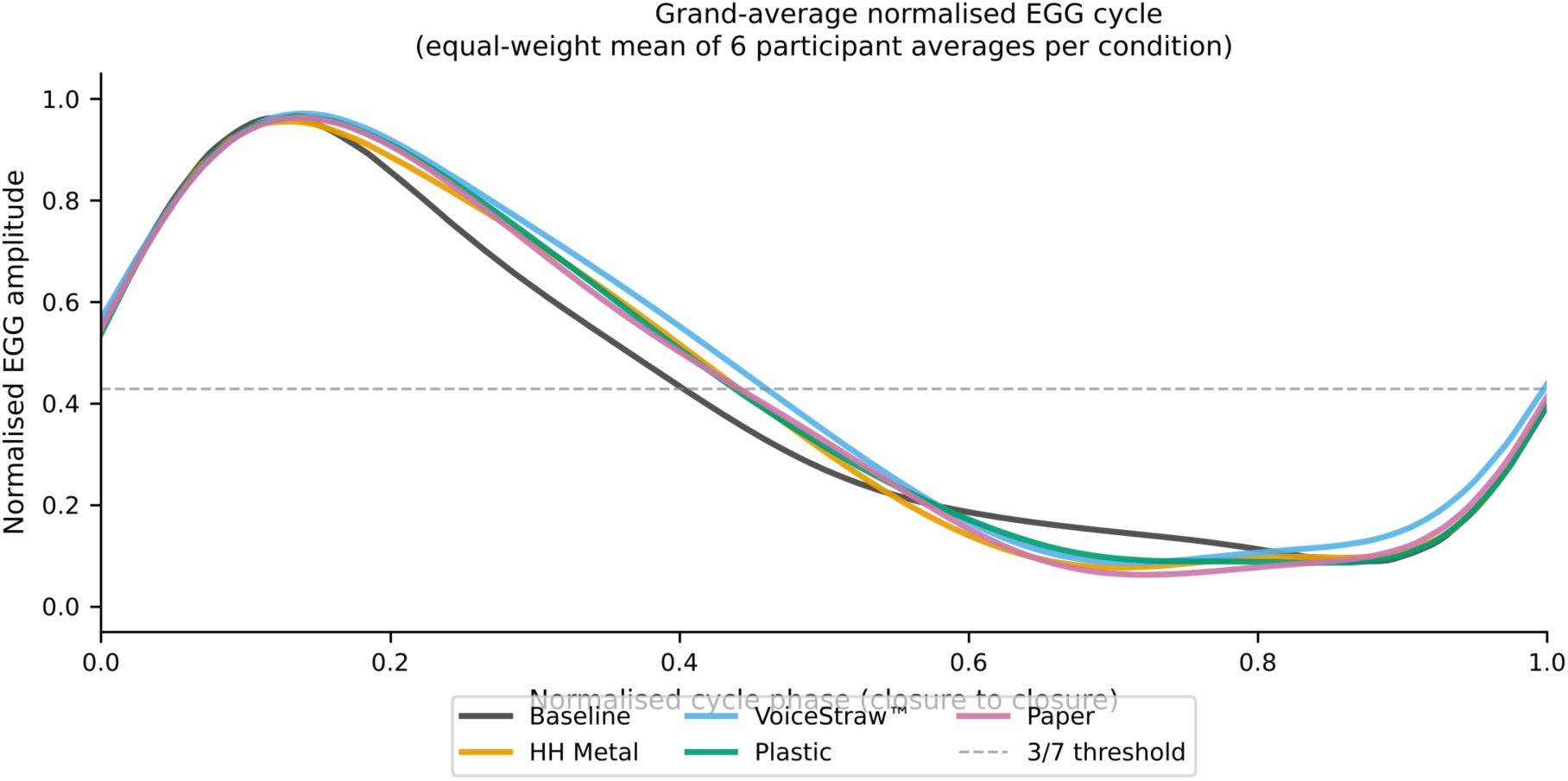
Grand-average normalised EGG cycle waveform for baseline and the four post-exercise straw conditions. Each curve is the equal-weight mean of six participant-specific mean cycles, each cycle amplitude-normalised to [0, 1] and resampled to 201 equally spaced phase points.

### 3.7 Overall descriptive pattern

Across the six participants, all post-exercise conditions showed higher mean CQ and lower waveform skewness than baseline. Mean rise/fall ratios were below baseline for the plastic condition; the Household metal, Metal VoiceStrawTM, and paper group means were marginally above baseline owing to high inter-individual variability, though 4–5 of 6 participants individually showed rf_ratio decreases for those conditions. Metal VoiceStrawTM showed the largest mean CQ increase and skewness decrease, while plastic showed the lowest mean rise/fall ratio. These descriptive patterns varied among participants and do not establish straw superiority. Table 3 summarizes group-level EGG measures. The dEGG skew column in Table 3 shows that derivative skewness was positive in most conditions (range 0.660–1.016), indicating greater skewness magnitude in the fall (opening) portion of the dEGG signal than in the rise (closing) portion. This is consistent with the NDAI direction reported in Section 3.5, providing a complementary confirmation of the asymmetry pattern.

**Table 3.** Group-level EGG measures. Values are mean ± SD across six participants. Cycle counts are retained complete cycles per 1-s recording.

| Condition | F0 (Hz) | CQ | OQ | Wave skew | Rise (ms) | Fall (ms) | Rise/Fall | dEGG skew | Cycles |
| --- | --- | --- | --- | --- | --- | --- | --- | --- | --- |
| Baseline | 280.3 ± 56.7 | 0.378 ± 0.112 | 0.622 ± 0.112 | 0.394 ± 0.208 | 0.476 ± 0.089 | 0.870 ± 0.298 | 0.588 ± 0.135 | 1.016 ± 0.434 | 280.8 ± 59.6 |
| Household metal | 260.9 ± 56.0 | 0.408 ± 0.091 | 0.592 ± 0.091 | 0.355 ± 0.316 | 0.558 ± 0.181 | 1.043 ± 0.374 | 0.600 ± 0.264 | 0.672 ± 0.541 | 257.7 ± 53.9 |
| Metal VoiceStraw™ | 275.4 ± 29.6 | 0.436 ± 0.085 | 0.564 ± 0.085 | 0.198 ± 0.236 | 0.554 ± 0.136 | 1.050 ± 0.381 | 0.628 ± 0.347 | 0.660 ± 0.651 | 272.3 ± 27.2 |
| Plastic | 271.8 ± 51.1 | 0.421 ± 0.066 | 0.579 ± 0.066 | 0.291 ± 0.233 | 0.573 ± 0.173 | 1.191 ± 0.536 | 0.562 ± 0.199 | 0.833 ± 0.532 | 251.8 ± 71.4 |
| Paper | 262.0 ± 52.9 | 0.405 ± 0.094 | 0.595 ± 0.094 | 0.333 ± 0.291 | 0.614 ± 0.260 | 1.042 ± 0.556 | 0.664 ± 0.282 | 0.727 ± 0.780 | 259.5 ± 52.5 |

### 3.8 EGG Spectral Metrics

Before presenting the group-level spectral metrics, participant F6 is shown as an illustrative example of the frequency-domain content of the EGG contact waveform. Figure 11 displays the normalised Welch power spectrum for each condition, demonstrating the harmonic structure of the EGG signal and condition-related differences in spectral shape. Figure 12 shows EGG spectrograms for each condition, illustrating the time-course stability of fundamental frequency and harmonic organisation across the 1-s recording window. These illustrative spectra and spectrograms are contact-waveform displays and should not be interpreted as acoustic microphone recordings.

**Figure 11.**
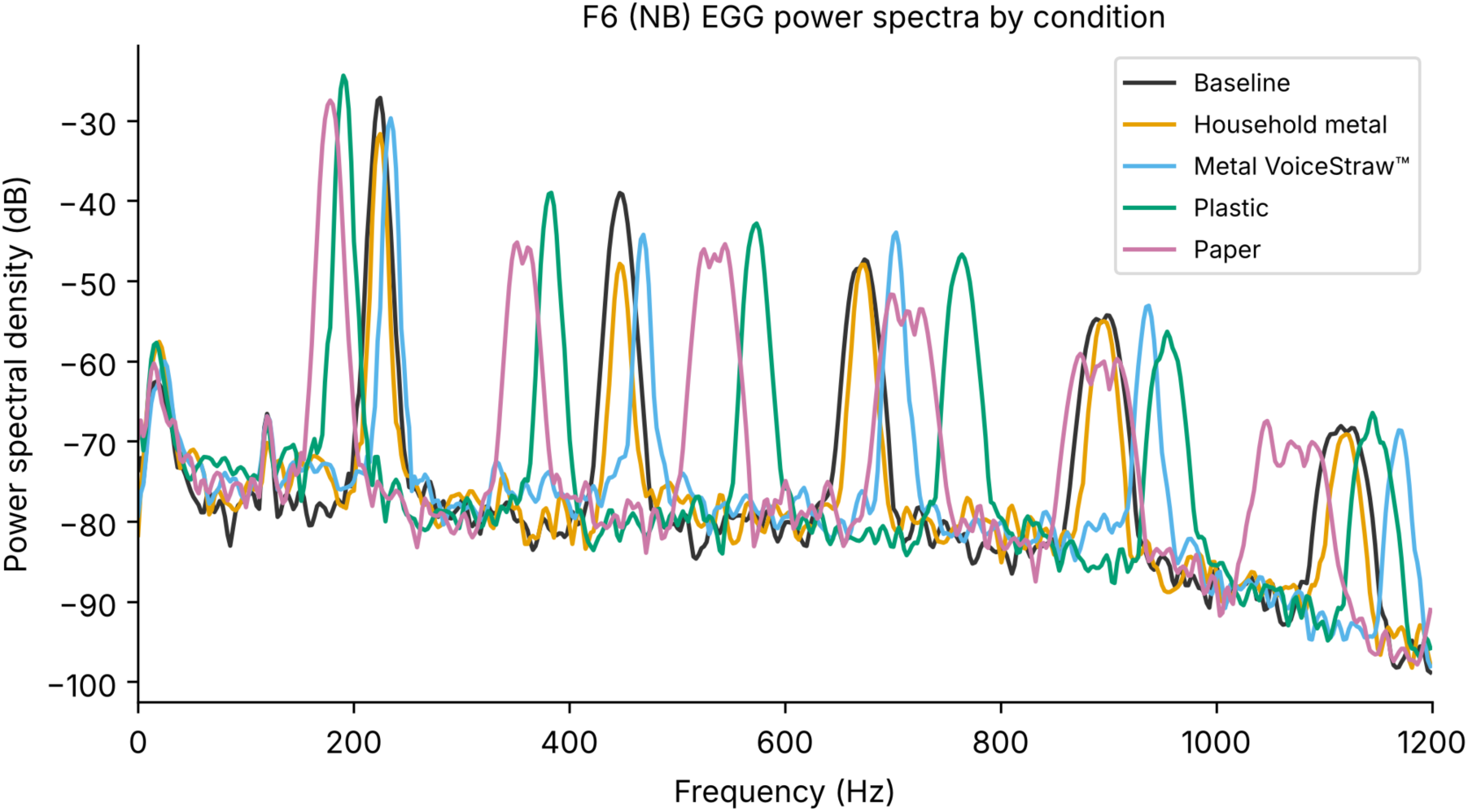
Illustrative normalised EGG power spectra for participant F6 under baseline, Household metal, Metal VoiceStraw™, plastic, and paper conditions. Spectra are computed via Welch’s method (Hann window, 2048-sample segments, 50% overlap, 4096-pt FFT) and displayed from 0 to 1000 Hz on a dB scale relative to the maximum harmonic peak to emphasise spectral shape. The regular spectral peaks correspond to harmonics of the EGG contact waveform.

**Figure 12.**
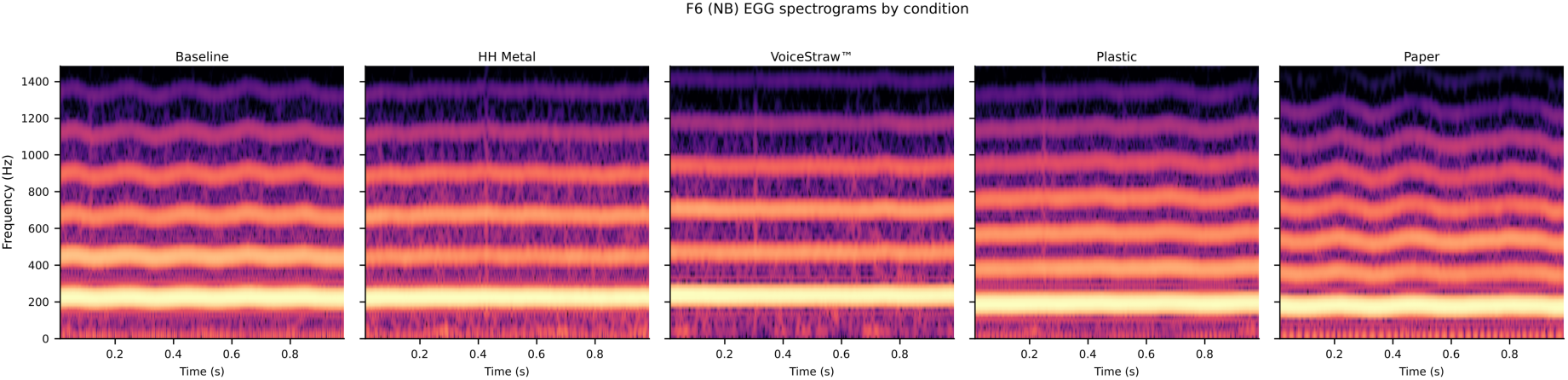
Illustrative EGG spectrograms for participant F6 under baseline, Household metal, Metal VoiceStraw™, plastic, and paper conditions. Each panel shows the time-varying spectral content of the EGG contact waveform over the full 1-s recording (0–1000 Hz, relative-dB scale, magma colormap). These are EGG contact-waveform spectrograms and should not be interpreted as acoustic microphone spectrograms.

With this illustrative context, the group-level analysis showed that EGG spectra retained a clearly harmonic organization in all five conditions, while the relative distribution of spectral power changed after the SOVT exercises. Group-mean spectral centroid decreased from 323.6 ± 55.9 Hz at baseline to 272.5 ± 58.0 Hz for Household metal, 287.8 ± 21.4 Hz for Metal VoiceStrawTM, 284.1 ± 56.8 Hz for plastic, and 281.3 ± 67.1 Hz for paper. Relative to each participant’s own baseline, the mean centroid changes were −15.3%, −9.6%, −11.8%, and −13.7%, respectively (Figure 13). Thus, descriptively, the post-SOVT EGG spectra showed a shift of the power-weighted frequency distribution toward lower frequencies. Group-level spectral metrics are summarized in Table 4.

**Figure 13.**
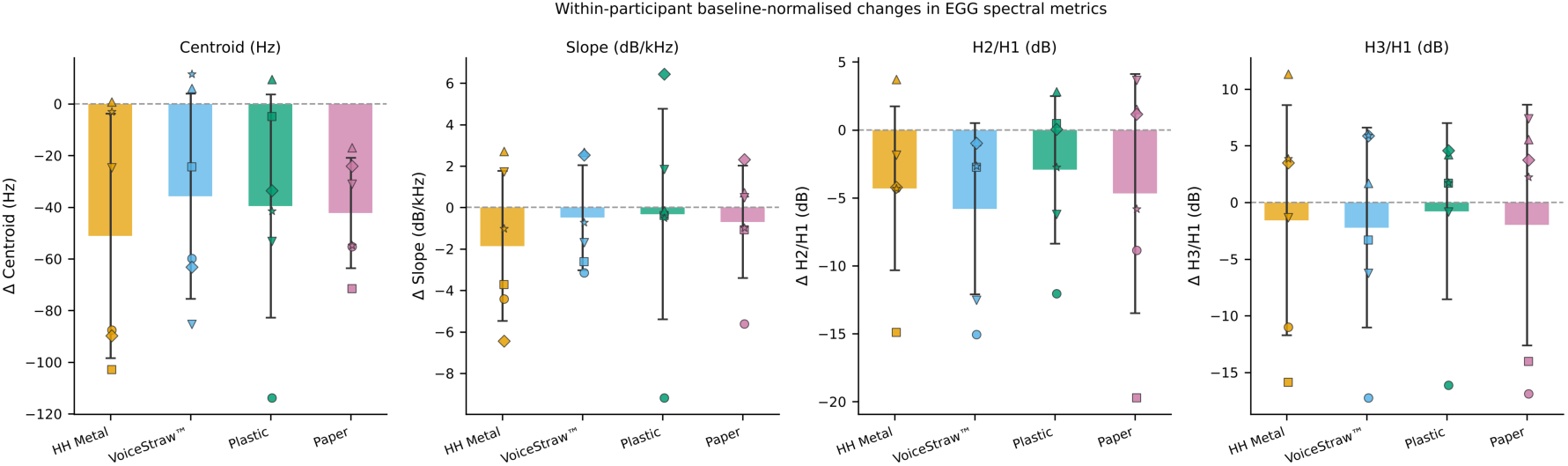
Group-mean baseline-normalised changes in EGG spectral centroid, spectral slope, H2/H1, and H3/H1 after the four SOVT straw conditions. Bars show mean ± SD across six participants; filled points show individual participant changes. Zero represents each participant’s own baseline value. Centroid is expressed as percentage change; spectral slope and harmonic ratios are arithmetic differences from baseline (dB).

**Table 4.** Group-level EGG spectral metrics by condition (mean ± SD).

| Condition | Centroid (Hz) | Slope (dB/kHz) | H2/H1 (dB) | H3/H1 (dB) |
| --- | --- | --- | --- | --- |
| Baseline | $323.6 \pm 55.9$ | $-10.2 \pm 2.2$ | $-10.3 \pm 2.6$ | $-20.6 \pm 6.3$ |
| Household metal | $272.5 \pm 58.0$ | $-12.0 \pm 3.5$ | $-14.6 \pm 6.1$ | $-22.2 \pm 9.0$ |
| Metal VoiceStraw™ | $287.8 \pm 21.4$ | $-10.7 \pm 2.1$ | $-16.1 \pm 4.4$ | $-22.8 \pm 5.3$ |
| Plastic | $284.1 \pm 56.8$ | $-10.5 \pm 4.7$ | $-13.2 \pm 3.6$ | $-21.4 \pm 3.2$ |
| Paper | $281.3 \pm 67.1$ | $-10.9 \pm 2.4$ | $-14.9 \pm 8.1$ | $-22.6 \pm 7.5$ |

H2/H1 also became more negative after all four straw conditions. The group mean was −10.3 dB at baseline and −14.6, −16.1, −13.2, and −14.9 dB after Household metal, Metal VoiceStrawTM, plastic, and paper, respectively. H3/H1 showed a smaller, similarly negative mean shift.

Spectral slope was −10.2 ± 2.2 dB/kHz at baseline and remained negative in all post-SOVT conditions. The largest mean baseline-normalized slope change occurred for Household metal (−1.8 dB/kHz); the changes for Metal VoiceStrawTM, plastic, and paper were −0.5, −0.3, and −0.7 dB/kHz. Participant-level spectral responses were heterogeneous. These group-level shifts are therefore presented descriptively as candidate spectral outcomes for confirmation in a larger study.

### 3.9 Self-reported and perceptual questionnaire outcomes

Comparative rankings were available for all six participants. In the complete F2 questionnaire, singing was rated easier after all four conditions; perceived voice change was reported after Metal VoiceStrawTM, plastic, and paper (’easier, less effort’) but not after Household metal. F2 ranked Metal VoiceStrawTM first, plastic second, paper third, and Household metal fourth and selected Metal VoiceStrawTM as easiest overall and most likely to be used for future warm-ups.

First-choice rankings across the six participants were Metal VoiceStrawTM 4/6 (66.7%), paper 2/6 (33.3%), plastic 0/6, and Household metal 0/6. Thus, four participants selected Metal VoiceStrawTM and two selected paper as their first choice; neither plastic nor Household metal was ranked first by any participant. Mean ranks (1 = most effective, 4 = least effective) were 1.67 for Metal VoiceStrawTM, 2.17 for paper, 2.50 for plastic, and 3.67 for Household metal. These values describe preference only and should not be interpreted as efficacy estimates or inferential evidence of a condition effect. The comparative ranking results are summarized in Table 5.

**Table 5.**
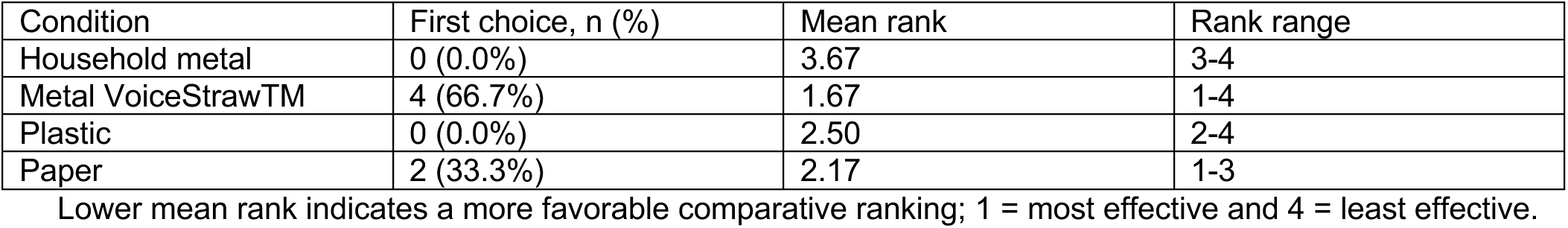
Comparative questionnaire ranking by straw condition (n = 6) Lower mean rank indicates a more favorable comparative ranking; 1 = most effective and 4 = least effective. These questionnaire findings are complementary perceptual data rather than evidence that one straw is superior.

| Condition | First choice, n (%) | Mean rank | Rank range |
| --- | --- | --- | --- |
| Household metal | 0 (0.0%) | 3.67 | 3-4 |
| Metal VoiceStraw™ | 4 (66.7%) | 1.67 | 1-4 |
| Plastic | 0 (0.0%) | 2.50 | 2-4 |
| Paper | 2 (33.3%) | 2.17 | 1-3 |

Open-ended responses described differences in effort, support, restriction, smoothness, resonance, fatigue, and ease of control, reinforcing inter-individual variability and the value of interpreting subjective preference alongside, rather than as a substitute for, EGG outcomes.

## 4. Discussion

### 4.1 Principal findings and interpretation

The principal contribution of this six-participant pilot is a reproducible multidimensional EGG framework for detecting immediate post-SOVT changes rather than identifying a superior straw. Across all four straw conditions, group means showed higher contact quotient (CQ), lower waveform skewness, lower rise/fall ratio, higher temporal skewness (s_t), and lower NDAI than the no-straw baseline. Considered together, the timing measures indicate a consistent descriptive shift toward a faster/steeper closing transition relative to the slower opening transition. The pattern was most evident for the Household metal, Metal VoiceStrawTM, and plastic conditions and smaller for paper. Participant-level trajectories nevertheless varied substantially, so these findings are hypothesis-generating rather than population-level condition effects.

The convergence of rise/fall timing, s_t, and NDAI is mechanistically relevant because rapid vocal-fold closing can contribute to rapid airflow declination. Prior aerodynamic work has linked flow skewing and rapid flow deceleration with glottal efficiency [8–10]. The present EGG findings are therefore consistent with the possibility that straw phonation promotes a closing pattern favorable to increased maximum flow declination rate. However, the observed spectral shifts (lower centroid, more negative H2/H1) are directionally opposite to what sharper closing would predict if F0 were held constant; the concurrent F0 rise may partly account for this discrepancy, but a direct test requires synchronized inverse-filtered airflow measurements. This interpretation remains indirect: EGG measures contact-related conductance rather than glottal airflow, glottal area, MFDR, or acoustic intensity. Simultaneous aerodynamic and acoustic measurements are needed to test the proposed mechanism directly.

Condition means were not identical. Metal VoiceStrawTM produced the largest descriptive CQ increase and waveform-skewness reduction, whereas plastic produced the lowest mean rise/fall ratio and highest mean s_t. These differences may reflect the acoustic/mechanical load created by straw geometry, but the present design cannot isolate material from internal diameter or other construction features. The strong inter-individual variability is also important: SOVT loading interacts with vocal-tract geometry, habitual adduction, pitch strategy, training history, and perception of back pressure. A larger study should therefore retain a within-subject repeated-measures design and model participant-specific responses.

### 4.2 Relationship to previous SOVT studies

The present results fit a literature in which post-SOVT responses depend on exercise type, tube geometry, dose, measurement timing, population, and outcome. Dargin and Searl [14] obtained aerodynamic and EGG measurements before and immediately after straw phonation, lip trill, and tongue trill in four singers; mean airflow, sound-pressure level, and EGG closed quotient tended to increase after the exercises, broadly consistent with the higher post-straw CQ observed here. However, CQ direction is not uniform across studies. Guzman et al. [21] reported decreased CQ during and after resonance-tube and narrow-straw phonation in a single intensively studied singer, and Mills et al. [22] reported a decreasing CQ trend after longer-duration straw tasks in 20 normal-voiced participants. These differences do not necessarily conflict: CQ is sensitive to task duration, pitch, loudness, tube geometry, timing, and the EGG algorithm used.

Tube geometry is particularly relevant to the present four-straw comparison. Mills et al. [22] manipulated tube length and showed duration-dependent post-exercise changes, while Smith and Titze [3] demonstrated that internal diameter strongly influences the pressure-flow behavior of flow-resistant tubes. Andrade et al. [11] further showed that different SOVT gestures are not physiologically interchangeable. Thus, the observed condition differences should be interpreted as responses to the complete straw condition rather than to material alone, especially because the four straws differed modestly in internal diameter.

To quantify whether back-pressure differences among the four straws could explain the observed EGG outcomes, flow resistance was estimated from the Hagen–Poiseuille approximation R = 8ηL/(πr⁴), using air dynamic viscosity η = 1.81 × 10⁻⁵ Pa·s and the nominal straw dimensions. Estimated resistances were 1.27, 1.99, 2.34, and 3.61 cmH₂O·s/L for the Metal VoiceStraw™, Paper, Plastic, and Household metal straws, respectively. All four values fall within the therapeutic range of approximately 1–5 cmH₂O·s/L identified by Smith and Titze [3] for flow-resistant tubes used in SOVT exercise, and at a representative expiratory airflow of 150 mL/s the estimated back-pressures ranged from 0.19 to 0.54 cmH₂O — less than 10 % of typical subglottal phonation pressure. Reynolds numbers at that flow rate ranged from approximately 2200 to 2900, indicating transitional rather than strictly laminar conditions, so the HP values slightly underestimate true resistance; the order-of-magnitude conclusion is nonetheless robust. Crucially, the rank ordering of resistance (Metal VoiceStraw™ lowest; Household metal highest) is inversely related to the rank ordering of post-exercise CQ change (Metal VoiceStraw™ largest change; Household metal among the smaller changes; Spearman ρ = −0.40). If back-pressure load were the primary mechanism driving glottal-contact increases, a positive resistance–outcome correlation would be expected; the negative value is inconsistent with that explanation. The condition differences therefore appear to arise from straw-specific factors other than resistance magnitude — most plausibly material vibration, surface texture, or subtle geometry effects. Direct measurement of each straw’s acoustic impedance and intraoral pressure in a larger study is nevertheless warranted (see Section 4.4).

The multidimensional findings are also consistent with studies showing that SOVT responses can appear in different measurement domains. Savareh et al. [24] reported changes in extrinsic laryngeal muscle activity and acoustic variables after an SOVT warm-up, while Ford et al. [25] found immediate acoustic, auditory-perceptual, and self-perceptual responses that varied among participants. Pozzali et al. [15] concluded that SOVT evidence remains heterogeneous across outcomes, and Meerschman et al. [26] evaluated straw phonation as one member of a broader SOVT therapy family. This literature supports the present cautious interpretation: measurable immediate changes are plausible, but a universal physiological response or superiority of one straw cannot be inferred from this pilot. Selected comparisons are summarized in Table 6.

**Table 6.**
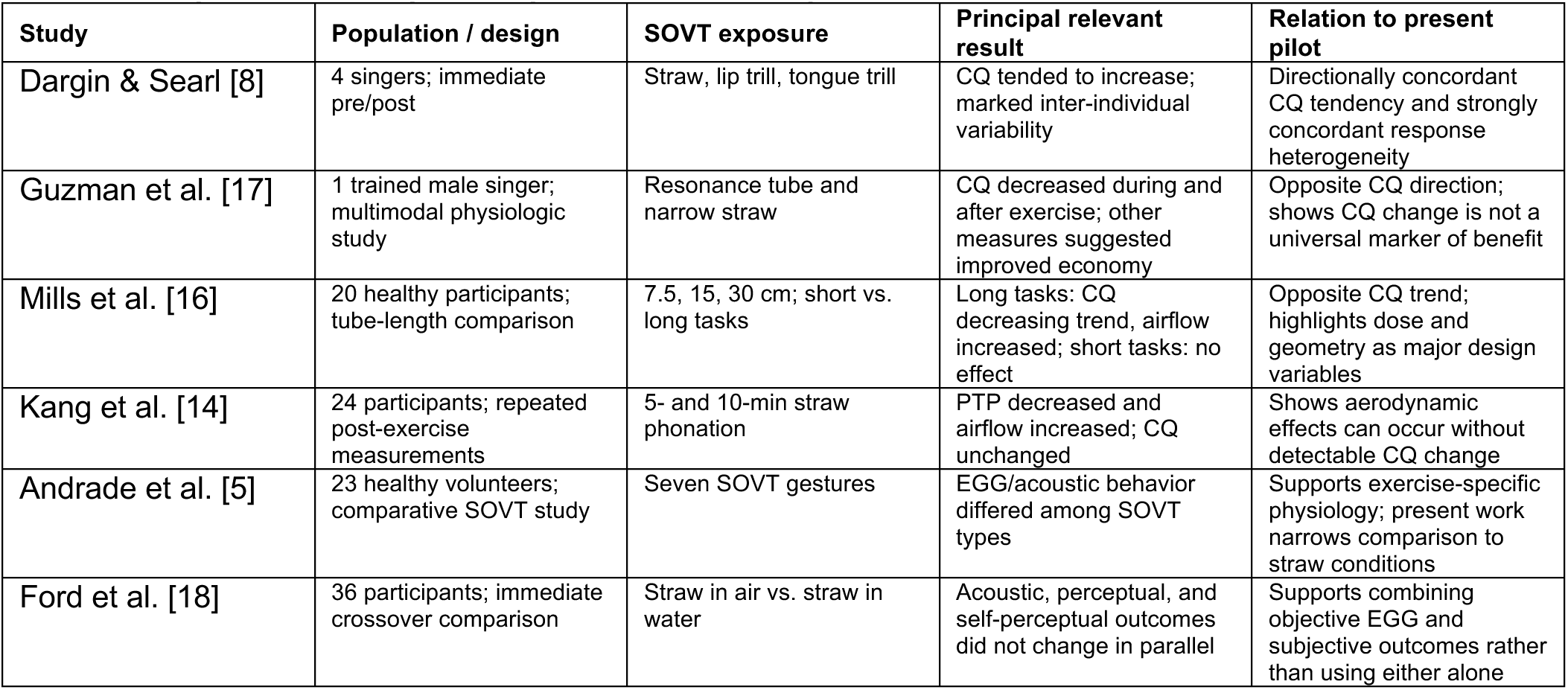
Comparison of the present pilot with selected prior SOVT studies.

The EGG spectral results provide a complementary frequency-domain description of the contact waveform. Post-SOVT group means showed lower spectral centroid and more negative H2/H1 than baseline while low-order harmonic organization was largely preserved. Because changes in the sharpness and temporal distribution of contact alter EGG harmonic composition, these spectral shifts are compatible with altered waveform morphology. They should not be interpreted as acoustic voice spectra or direct measures of voice quality. Their value here is as candidate EGG endpoints to be tested against synchronized acoustic and aerodynamic measures in a larger study.

The continuous microphone recording had a deliberately supporting role in this methodology. It provided an independent record of protocol delivery and session chronology that could be consulted when organizing EGG files, confirming that the intended brief phonation occurred, distinguishing test events from warm-up and rest periods, and identifying obvious interruptions or task deviations. This type of procedural audio record is useful even when acoustic outcomes are outside the study aims because it improves traceability and quality assurance without requiring the audio to be interpreted as a quantitative voice dataset.

The frequency-domain analysis provides a complementary view of the same EGG contact waveform. Taken together, this pattern indicates redistribution of EGG spectral power toward the lower-frequency portion of the contact waveform spectrum without a consistent loss of harmonic organization. The effect should not be described as an acoustic “brighter/darker” voice change because these are EGG spectra, not microphone spectra.

The spectral findings are also relevant to the closure-asymmetry interpretation developed from the supporting rise/fall timing measure and the principal temporal-skewness and NDAI outcomes. A change in the sharpness and temporal distribution of vocal-fold contact necessarily changes the harmonic composition of the EGG waveform; therefore, the observed spectral shifts provide an independent frequency-domain descriptor of altered contact-waveform shape. However, the present data do not establish a one-to-one mapping between EGG H2/H1 or spectral slope and MFDR. MFDR is an airflow-derived measure, and the proposed link between faster EGG contact transition, altered EGG spectral structure, and increased airflow declination must be tested directly with synchronized inverse-filtered flow or other validated glottal-flow measurements.

The directionally similar centroid and H2/H1 shifts across condition means identify potentially useful candidate endpoints, but participant-level variability and the inter-individual variability in baseline H2/H1 (SD ± 2.6 dB) show why confirmation in a larger repeated-measures sample is required. A future adequately sized protocol should prespecify the spectral frequency range and harmonic-search rule, collect repeated baseline recordings, and test whether these EGG spectral measures covary with CQ, s_t, NDAI, and independent acoustic or aerodynamic outcomes.

### 4.3 New contribution of the methodology

The methodological contribution is the integration of conventional and complementary EGG measures within one fixed pipeline. Complete cycles were identified from dEGG closure landmarks, cycle duration and amplitude were normalized, CQ/OQ were calculated with an explicitly defined opening criterion, and waveform shape was characterized using rise/fall timing, s_t, NDAI, average-cycle morphology, and spectral descriptors. The 35%, 3/7 (42.9%), and 50% opening-threshold sensitivity analysis showed that absolute CQ and asymmetry values depend on the criterion, as expected, but the principal baseline-to-straw directions were preserved. This supports the robustness of the descriptive pattern while emphasizing that EGG quotients are method dependent.

The framework also identifies variables that should be controlled in a larger study. Exercise duration, pitch range, loudness, posture, hydration, time from exercise cessation to recording, electrode placement, and recording level should be standardized or documented. Repeated baseline recordings would help quantify measurement repeatability. The microphone recording remains useful as procedural documentation for confirming task execution, chronology, and interruptions even when microphone-derived acoustic outcomes are not analyzed.

### 4.4 Implications for a larger confirmatory study

A larger confirmatory study should prespecify a small number of primary outcomes, particularly temporal asymmetry measures and CQ, and treat the remaining waveform and spectral variables as secondary or mechanistic endpoints. Sample size should be based on a clinically or physiologically meaningful difference and a conservative estimate of within-participant variability rather than the largest change observed in this pilot. With an adequately sized sample, repeated-measures mixed-effects models could separate participant variability from condition-related differences and accommodate missing observations.

Future experiments should separate geometry from material by matching straw length and internal diameter as closely as possible and directly measuring each straw’s acoustic/flow resistance. Simultaneous airflow, oral pressure, calibrated acoustic intensity, and EGG would allow direct testing of whether the observed steeper EGG closing pattern corresponds to increased MFDR, glottal efficiency, or spectral excitation. The concealment holder should be retained because it reduces visual expectancy bias in subjective comparisons, although it does not provide complete blinding because tactile and airflow cues may remain.

### 4.5 Limitations

The study has several limitations. The sample of six female community singers is too small for population-level inference or conclusions about superiority among straw conditions. Post-exercise F0 was not identical across conditions, and the four straws differed in geometry as well as material. EGG is a relative contact measure affected by electrode placement, neck anatomy, preprocessing, polarity, and landmark definition; although threshold sensitivity testing supported the directional robustness of the findings, it does not make the opening landmark anatomically exact. The 1-s recordings also provide only brief observations of each condition.

The baseline-first design means that every post-straw recording follows at least one prior phonation condition; warm-up effects, F0 drift, and electrode acclimatization cannot be separated from straw-specific effects. The 10-minute washout interval may be insufficient: Kang et al. [23] reported lingering aerodynamic and EGG effects of straw phonation beyond 10 minutes. These are irreducible confounds of the present design that should be addressed in the confirmatory study by including repeated baselines and a matched non-SOVT phonation control.

The microphone recordings were retained for procedural documentation rather than acoustic analysis, so no claims about acoustic voice changes are made from those recordings. Likewise, EGG spectral magnitude and harmonic ratios are placement-and processing-dependent and should be interpreted as within-participant descriptors of contact-waveform shape rather than direct acoustic measures. Finally, the holder concealed straw appearance but could not eliminate tactile or airflow cues and therefore should not be described as complete participant blinding.

## 5. Conclusions

This pilot study demonstrates a feasible multidimensional methodology for quantifying immediate changes after straw-phonation SOVT exercises using EGG cycle analysis together with participant-reported experience. Across six participants, all four post-exercise straw conditions showed the same descriptive group-level directions for several measures: higher CQ, lower waveform skewness, a lower supporting rise/fall time ratio, greater temporal asymmetry by the principal s_t measure, and lower derivative asymmetry by NDAI relative to baseline. The magnitude of these changes differed by condition and by participant; these differences are reported descriptively and are not treated as population-level material-specific effects. Accordingly, the principal new finding is not that one straw material is superior, but that the proposed analysis pipeline can detect multiple, physiologically interpretable dimensions of post-SOVT waveform change while preserving the substantial individual variability that characterizes singer responses. The contemporaneous microphone recording served only as procedural documentation and EGG quality-control support and was not analyzed as an acoustic outcome. A particularly important hypothesis-generating result is the coordinated post-SOVT shift toward shorter EGG rise time, lower rise/fall ratio, higher s_t, and lower NDAI. The Household-metal, Metal VoiceStrawTM, and plastic conditions showed the clearest combined shifts, while paper showed a smaller change in the same group-mean direction. This pattern is consistent with a faster late-closing/contact transition and is mechanistically relevant because rapid glottal-area closure and vocal-tract inertance are established contributors to MFDR, which is linked to acoustic excitation and vocal intensity [7–10]. The study does not directly demonstrate increased MFDR or efficiency; rather, it identifies EGG asymmetry measures as promising noninvasive markers to be validated against simultaneous airflow, pressure, acoustic, and glottal-area measurements in the next-stage study. These spectral measures complement the time-domain asymmetry measures and provide additional candidate endpoints for a larger multimodal study, but they should not be interpreted as direct acoustic spectra or as direct measurements of MFDR. Reanalysis with 35%, 3/7, and 50% opening thresholds preserved the direction of the CQ, temporal-asymmetry, and NDAI changes, indicating that the central descriptive pattern was robust to reasonable criterion-level variation.

Comparison with prior work shows that the present findings occupy a plausible but not universal part of the SOVT response spectrum. The higher descriptive CQ is directionally consistent with Dargin and Searl [14], whereas Guzman et al. [21] and Mills et al. [22] reported lower/decreasing CQ and Kang et al. [23] found no significant CQ change despite aerodynamic improvement. Rather than constituting a simple contradiction, this pattern indicates that CQ is sensitive to exercise dose, straw geometry, population, recording timing, and analysis method and should not be used alone as a marker of SOVT benefit. The non-identical questionnaire preferences and perceived changes are similarly consistent with Ford et al. [25], who showed that objective and self-perceptual outcomes need not change in parallel. A larger, adequately powered, randomized/counterbalanced repeated-measures study with tightly characterized straw geometry and material, standardized exercise dose, synchronized EGG and, where specifically required by the research question, dedicated acoustic/aerodynamic measurements, repeated baselines, and predefined primary outcomes is therefore warranted. Such a study can determine whether the descriptive differences observed here represent reproducible material-related effects, interactions with geometry and dose, or participant-specific responses.

## Institutional Review Board Statement

The study protocol (#2025-0933) was approved by the University of Cincinnati Institutional Review Board.

## Informed Consent Statement

Written informed consent was obtained from participants before study-specific procedures.

## Author Contributions

M.L. recruited participants, conducted the tests, analyzed the data, and prepared the manuscript. L.O. initiated the study, provided the experimental facility and software, managed the project, and reviewed the results and manuscript.

## Data Availability Statement

The raw EGG data (30 TDMS files) and analysis code supporting this study will be deposited in a public repository (Zenodo or OSF) upon acceptance. The DOI will be inserted here prior to publication. Data are currently available from the corresponding author upon reasonable request.

## References

1. Titze, I.R. Voice training and therapy with a semi-occluded vocal tract: Rationale and scientific underpinnings. J. Speech Lang. Hear. Res. 2006, 49, 448–459. 10.1044/1092-4388(2006/035).

2. Maxfield, L.; Titze, I.; Hunter, E.; Kapsner-Smith, M. Intraoral pressures produced by thirteen semi-occluded vocal tract gestures. Logoped. Phoniatr. Vocol. 2015, 40, 86–92. 10.3109/14015439.2014.913074.

3. Smith, S.L.; Titze, I.R. Characterization of flow-resistant tubes used for semi-occluded vocal tract voice training and therapy. J. Voice 2017, 31, 113.e1–113.e8. 10.1016/j.jvoice.2016.04.001.

4. Titze, I.R. Phonation threshold pressure measurement with a semi-occluded vocal tract. J. Speech Lang. Hear. Res. 2009, 52. 10.1044/1092-4388(2009/08-0110).

5. Näger, C.; Kniesburges, S.; Tur, B.; Schoder, S.; Becker, S. An investigation of acoustic back-coupling in human phonation on a synthetic larynx model. Bioengineering 2023, 10, 1343. 10.3390/bioengineering10121343.

6. Michaud-Dorko, J.; Farbos de Luzan, C.; Dion, G.R.; Gutmark, E.; Oren, L. Comparison of aerodynamic and elastic properties in tissue and synthetic models of vocal fold vibrations. Bioengineering 2024, 11, 834. 10.3390/bioengineering11080834.

7. Gauffin, J.; Sundberg, J. Spectral correlates of glottal voice source waveform characteristics. J. Speech Hear. Res. 1989, 32, 556–565. 10.1044/jshr.3203.556.

8. Titze, I.R. Theoretical analysis of maximum flow declination rate versus maximum area declination rate in phonation. J. Speech Lang. Hear. Res. 2006, 49, 439–447. 10.1044/1092-4388(2006/034).

9. Titze, I.R.; Laukkanen, A.-M. Can vocal economy in phonation be increased with an artificially lengthened vocal tract? A computer modeling study. Logoped. Phoniatr. Vocol. 2007, 32, 147–156. 10.1080/14015430701439765.

10. Oren, L.; Khosla, S.; Dembinski, D.; Ying, J.; Gutmark, E. Direct measurement of planar flow rate in an excised canine larynx model. Laryngoscope 2015, 125, 383–388. 10.1002/lary.24866.

11. Andrade, P.A.; Wood, G.; Ratcliffe, P.; Epstein, R.; Pijper, A.; Švec, J.G. Electroglottographic study of seven semi-occluded exercises: LaxVox, straw, lip-trill, tongue-trill, humming, hand-over-mouth, and tongue-trill combined with hand-over-mouth. J. Voice 2014, 28, 589–595. 10.1016/j.jvoice.2013.11.004.

12. Titze, I.R.; Hunter, E.J. Feasibility of measurement of a voice range profile with a semi-occluded vocal tract. Logoped. Phoniatr. Vocol. 2011, 36, 32–39. 10.3109/14015439.2010.548828.

13. Titze, I.R.; Maxfield, L.; Titze-Cox, K. Optimizing diameter, length, and water immersion in flow-resistant tube vocalization. J. Voice 2025, 39, 403–409. 10.1016/j.jvoice.2022.09.029.

14. Dargin, T.C.; Searl, J. Semi-occluded vocal tract exercises: Aerodynamic and electroglottographic measurements in singers. J. Voice 2015, 29, 155–164. 10.1016/j.jvoice.2014.05.009.

15. Pozzali, I.; Pizzorni, N.; Ruggeri, A.; Schindler, A.; Dal Farra, F. Effectiveness of semi-occluded vocal tract exercises (SOVTEs) in patients with dysphonia: A systematic review and meta-analysis. J. Voice 2024, 38, 245.e17–245.e35. 10.1016/j.jvoice.2021.06.009.

16. Riquelme-Muñoz, C.; Labarca-Farias, A.; Contreras-Arancibia, C. Effectiveness of vocal exercises in singers: A scoping review. J. Voice 2025. 10.1016/j.jvoice.2025.09.029.

17. Jiang, W.; Zheng, X.; Farbos de Luzan, C.; Oren, L.; Gutmark, E.; Xue, Q. The effects of negative pressure induced by flow separation vortices on vocal fold dynamics during voice production. Bioengineering 2023, 10, 1215. 10.3390/bioengineering10101215.

18. Howard, D.M. Variation of electrolaryngographically derived closed quotient for trained and untrained adult female singers. J. Voice 1995, 9, 163–172. 10.1016/S0892-1997(05)80250-4.

19. Abramson, A.S.; Tiede, M.K.; Luangthongkum, T. Voice register in Mon: Acoustics and electroglottography. Phonetica 2015, 72, 237–256. 10.1159/000441728.

20. Rothenberg, M.; Mahshie, J.J. Monitoring vocal fold abduction through vocal fold contact area. J. Speech Hear. Res. 1988, 31, 338–351. 10.1044/jshr.3103.338.

21. Guzman, M.; Laukkanen, A.-M.; Krupa, P.; Horáček, J.; Švec, J.G.; Geneid, A. Vocal tract and glottal function during and after vocal exercising with resonance tube and straw. J. Voice 2013, 27, 523.e19–523.e34. 10.1016/j.jvoice.2013.02.007.

22. Mills, R.D.; Rivedal, S.; DeMorett, C.; Maples, G.; Jiang, J.J. Effects of straw phonation through tubes of varied lengths on sustained vowels in normal-voiced participants. J. Voice 2018, 32, 386.e21–386.e29. 10.1016/j.jvoice.2017.05.015.

23. Kang, J.; Xue, C.; Piotrowski, D.; Gong, T.; Zhang, Y.; Jiang, J.J. Lingering effects of straw phonation exercises on aerodynamic, electroglottographic, and acoustic parameters. J. Voice 2019, 33, 810.e5–810.e11. 10.1016/j.jvoice.2018.05.002.

24. Savareh, S.; Moradi, N.; Shaterzadeh Yazdi, M.J.; Soltani, M.; Latifi, M. Immediate effects of semi-occluded vocal tract exercises as a vocal warm-up in singers. J. Voice 2023, 37, 875–880. 10.1016/j.jvoice.2021.05.014.

25. Ford, D.S.; Hunter, E.J.; Deliyski, D.D. Immediate effects of semi-occluded vocal tract exercises on acoustic, auditory-perceptual, and self-perceptual measures of voice production. Folia Phoniatr. Logop. 2024, 76, 467–481. 10.1159/000536002.

26. Meerschman, I.; Van Lierde, K.; Ketels, J.; Coppieters, C.; Claeys, S.; D’haeseleer, E. Effect of three semi-occluded vocal tract therapy programmes on the phonation of patients with dysphonia: Lip trill, water-resistance therapy and straw phonation. Int. J. Lang. Commun. Disord. 2019, 54, 50–61. 10.1111/1460-6984.12431.

